# A cytomegalovirus immunevasin triggers integrated stress response-dependent reorganization of the endoplasmic reticulum

**DOI:** 10.1101/641068

**Authors:** Hongbo Zhang, Clarissa Read, Christopher C. Nguyen, Mohammed N.A. Siddiquey, Chaowei Shang, Cameron M. Hall, Jens von Einem, Jeremy P. Kamil

**Affiliations:** Department of Microbiology and Immunology, LSU Health Sciences Center, Shreveport, Louisiana, USA; Center for Molecular and Tumor Virology, LSU Health Sciences Center, Shreveport, Louisiana, USA; Research Core Facility, LSU Health Sciences Center, Shreveport, Louisiana, USA; Institute of Virology, Ulm University Medical Center, Ulm University Ulm, Germany; Central Facility for Electron Microscopy, Ulm University Ulm, Germany

## Abstract

Human cytomegalovirus (HCMV) encodes an ER-resident glycoprotein, UL148, which activates the unfolded protein response (UPR) but is fully dispensable for viral replication in cultured cells. Hence, its previously ascribed roles in immune evasion and modulation of viral cell tropism are hypothesized to cause ER stress. Here, we show that UL148 is necessary and sufficient to drive the formation of large ER-derived structures that occupy up to 7% of the infected cell cytosol. The structures are found to be sites where UL148 coalesces with cellular proteins involved in ER quality control, such as Hrd1 and EDEM1. Transmission electron microscopy analyses reveal the structures to be comprised of tortuous, densely packed and apparently collapsed ER membranes that connect to distended cisternae. During induced ectopic expression of UL148-GFP fusion protein, punctate signals traffic to accumulate at prominent structures that exhibit poor recovery of fluorescence after photobleaching. Small molecule blockade of the integrated stress response (ISR) prevents the formation of puncta, leading to a uniform reticular fluorescent signal. Accordingly, ISR inhibition during HCMV infection abolishes the coalescence of UL148 and Hrd1 into discrete structures, which argues that UL148 requires the ISR to cause ER reorganization. Given that UL148 stabilizes immature forms of a receptor binding subunit for a viral envelope glycoprotein complex important for HCMV infectivity, our results imply that stress-dependent ER remodeling contributes to viral cell tropism.

**IMPORTANCE:** Perturbations to ER morphology occur during infection with various intracellular pathogens and in certain genetic disorders. We identify that an HCMV gene product, UL148, profoundly reorganizes the ER during infection, and is sufficient to do so when expressed on its own. Our results reveal that UL148-dependent reorganization of the ER is a prominent feature of HCMV infected cells. Moreover, we find that this example of virally induced organelle remodeling requires the integrated stress response (ISR), a stress adaptation pathway that contributes to a number of disease states. Since ER reorganization accompanies the roles of UL148 in HCMV cell tropism and intracellular retention of the immune cell co-stimulatory ligand CD58, our results may have implications for understanding the mechanisms involved. Furthermore, our findings provide a basis to utilize UL148 as a tool to investigate organelle responses to stress and to identify novel drugs targeting the ISR.

## INTRODUCTION

UL148 is a human cytomegalovirus (HCMV) ER-resident glycoprotein that plays roles in evasion of cell-mediated immunity and shows intriguing effects on cell tropism. During infection of epithelial cells, viruses disrupted for *UL148* replicate to produce roughly 100-fold enhanced levels of infectious progeny virions compared to wildtype (1). These effects correlate with reduced expression of glycoprotein O (gO), a subunit of a heterotrimeric viral glycoprotein H (gH) / glycoprotein L (gL) complex (gH/gL/gO) that is required for the infectivity of cell-free virions (2–4). The presence of gO in the context of the heterotrimer endows the virus with the capacity to utilize the platelet derived growth factor receptor *α* (PDGFR*α*) as an entry receptor (5–7). Accordingly, UL148 has been found to stabilize immature forms of gO prior their assembly into gH/gL/gO heterotrimers (1, 8). Despite that UL148 does not stably associate with gO, the data suggest an interaction with gH (1).

UL148 also physically associates with CD58 (LFA-3), a co-stimulatory ligand for natural killer cells and T-lymphocytes, preventing its presentation at cell surface (9). Although the mechanisms by which UL148 stabilizes gO and retains CD58 within the ER remain unknown, UL148 strongly contributes to activation of the unfolded protein response (UPR) during infection, and is sufficient to activate the UPR when ectopically expressed in non-infected cells (10). UL148 co-purifies from infected cells with SEL1L, an adaptor subunit of ER-based E3 ubiquitin ligase Hrd1 that plays important roles in ER-associated degradation (ERAD) of terminally misfolded glycoproteins (8). This suggests a physical interaction with ERAD machinery, which may be germane to the mechanism by which UL148 activates the UPR.

Here, we show that the expression of UL148 is necessary and sufficient to induce unusual ER structures at which large quantities of ER factors involved in glycoprotein quality control accumulate. In electron microscopy analyses, we find that the UL148 induced ER-structures are comprised of densely packed, tortuous ER membranes that form connections with tubules of highly distended cisternal space. Furthermore, additional data from inhibitor studies strongly argue that ER remodeling triggered by UL148 requires the integrated stress response. Overall, our results reveal a striking ER perturbation induced by HCMV, and which is entirely controlled by a single viral gene product. These findings may have important implications for understanding how UL148 regulates viral cell tropism and/or contributes to viral evasion of cell-mediated immunity.

## RESULTS

### UL148 causes reorganization of ER quality control proteins into unusual globular structures

The HCMV ER-resident glycoprotein UL148 was previously observed to co-localize with the ER marker calnexin during infection (1). Nonetheless, calnexin staining did not show the uniform reticular pattern characteristic for the ER marker. We later noticed that cells infected with a *UL148*-null virus showed uniform calnexin staining (see below). To formally determine whether UL148 influences calnexin localization, we compared fibroblasts at four days post-infection with either wildtype strain TB40/E virus (TB_WT) or a UL148-null mutant (TB_148_STOP_)(8, 10), each derived from an infectious bacterial artificial chromosome (BAC) clone of HCMV strain TB40/E (11) (**FIG 1**). In cells infected with wildtype virus, calnexin antibodies stained unusual globular structures at the cell periphery, as expected (1) (**FIG 1A, 1C, SI FIG S1A**). However, in cells infected with the *UL148*-null virus, calnexin staining was uniform throughout the cytosol (**FIG 1B-C**), as would be expected for an ER marker in uninfected cells. The staining pattern for Hrd1, another ER marker, likewise indicated accumulation at unusual globular structures during wildtype HCMV infection, but not during infection with *UL148*-null mutant viruses (**FIG 1, S1A**). Because calnexin and Hrd1 staining showed uniform distribution in *UL148*-null virus infected cells, and because *UL148* is fully dispensable for efficient viral replication in fibroblasts (1), these results suggest that redistribution of these two ER markers depends on UL148.

**FIG 1.**
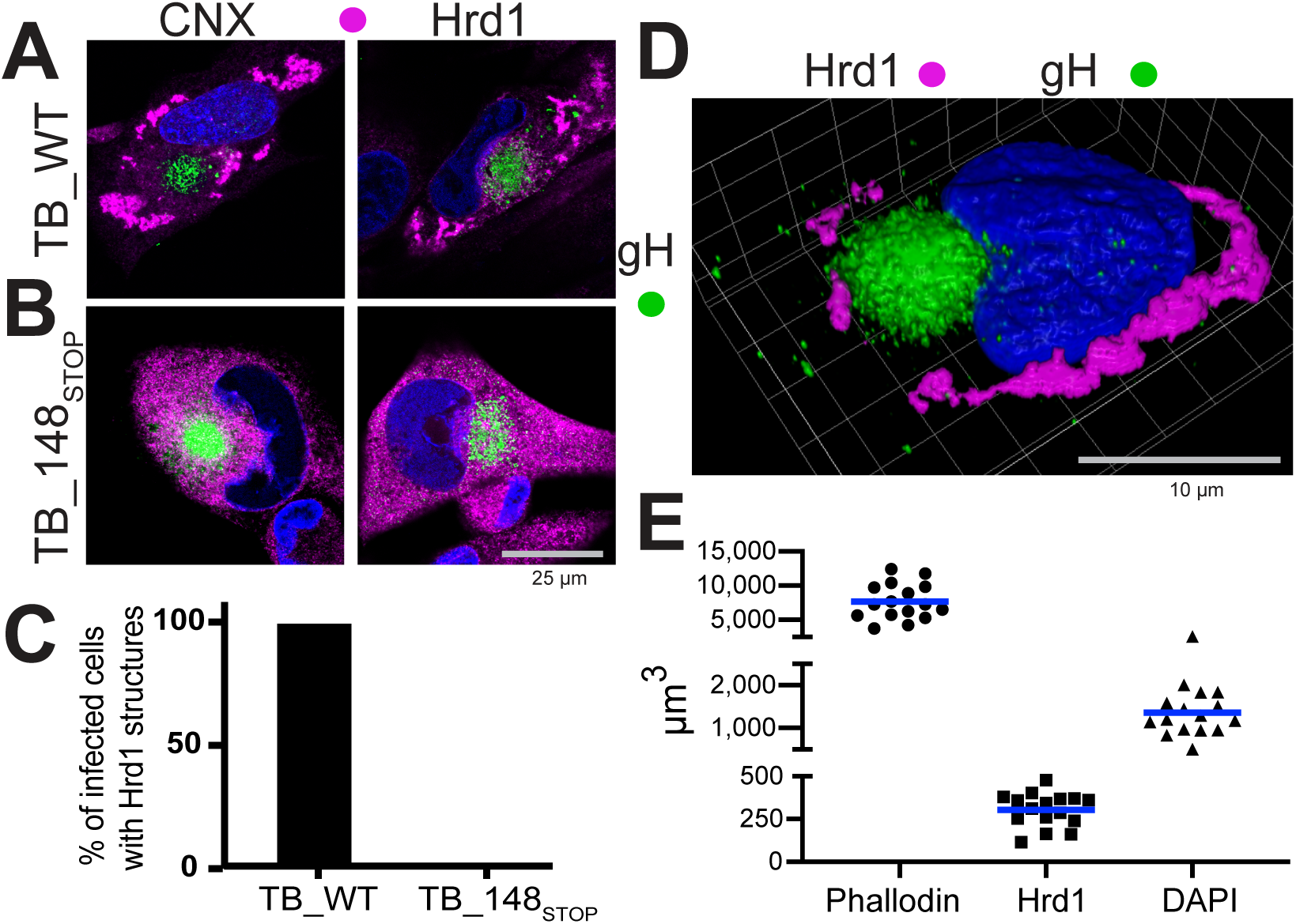
UL148 reorganizes ER markers into anomalous structures during HCMV infection. Fibroblasts infected at MOI 1 TCID_50_/cell with either (**A**) wildtype HCMV strain TB40/E (TB_WT) or (**B**) an isogenic *UL148*-null mutant (TB_148_STOP_), were fixed at 96 h postinfection (hpi) and imaged by confocal microscopy after staining with antibodies specific for calnexin or Hrd1 (magenta) and glycoprotein H (gH, green), as indicated. DAPI (blue) signal was used to counterstain nuclei. **(C)** Percentage of fibroblasts that contain Hrd1 structures at 96 hpi; fifty gH-positive cells for each condition were scored for the presence or absence of UL148 structures (also see SI Fig S1A). (**D**) 3D confocal image projection of a TB_WT infected fibroblast stained at 96 hpi for Hrd1 and gH. **(E)** Volumetric measurements were made from sixteen infected cells, stained with phalloidin-AlexaFluor 594 conjugate to estimate total cell volume, with Hrd1 antibody to estimate the volume of UL148-dependent ER structures, and with DAPI to estimate nuclear volume.

As expected (12, 13), antibodies specific for glycoprotein H (gH), a viral envelope glycoprotein, stained a juxtanuclear compartment, termed the cytoplasmic virion assembly compartment (cVAC), which does not involve the ER (**FIG 1**). Similarly contrasting staining patterns for Hrd1 and/or calnexin were observed in cells infected with wildtype versus *UL148*-null mutants of clinical HCMV strains Merlin and TR (**FIG S1**). Reciprocally, we restored a functional *UL148* at its native locus in the context of a BAC clone of HCMV strain AD169, which spontaneously lost most of the gene in the course of extensive genetic rearrangements that occurred during serial passage in cultured cells. In cells infected with AD169 repaired for UL148, but not the *UL148*-null parental virus, the Hrd1 staining pattern showed prominent globular structures (**FIG S1**).

Because similar differences in Hrd1 staining are observed between wildtype and *UL148*-null infections of THP-1 macrophages, ARPE-19 epithelial cells and fibroblasts (**FIG 1, FIG S1**), we concluded that UL148-dependent reorganization of ER markers occurs in multiple cell types. The calnexin staining pattern seen with four different primary clinical isolates from patient throat swabs likewise indicated punctate globular structures. These observations, taken together with results from formal comparisons of wildtype versus *UL148*-null mutants of four different BAC-cloned HCMV strains (**FIG 1, FIG S1**), argue that UL148 profoundly affects the ER during natural infection.

We measured the three-dimensional volume of UL148-dependent Hrd1 structures from sixteen cells fixed at 96 h post infection (hpi) with strain TB40/E. On average, the structures occupied 303.3 µm^3^ (SEM: +/-24.8) out of a total cell volume of 7657 µm^3^ (SEM:+/-651.0), which was estimated using a phalloidin-fluorophore conjugate to detect the actin cytoskeleton (**FIG 1D-E**). Subtracting the volume of nuclei, as indicated by 4′,6-diamidino-2-phenylindole [DAPI] (average nucleus: 1360 µm^3^, SEM:+/-129.4), we calculate that, on average, the structures occupy 5.2% of the cytosolic volume (SEM: +/-0.61%, range: 2.0% – 7.0%). Based on these findings, taken together with results comparing additional wildtype (WT) versus *UL148*-null mutant HCMV strains (**FIG S1**), we conclude that the UL148-dependent ER structures are a prominent feature of HCMV-infected cells.

UL148 substantially contributes to activation of the unfolded protein response (UPR) during HCMV infection (10), and co-purifies from infected cells with SEL1L (8), an adaptor subunit for the E3 ubiquitin ligase Hrd1, which plays crucial roles in ER-associated degradation (ERAD) of terminally misfolded glycoprotein substrates (14). Hence, the accumulation of Hrd1 and calnexin at unusual structures during wildtype but not *UL148*-null infection may suggest that structures form in response to defects in ER quality control (ERQC) caused by UL148. Under conditions of proteasome inhibition, overexpression of certain misfolded glycoproteins causes cellular factors involved in ERQC, such as calnexin, to compartmentalize from the rest of the ER, while other ER markers such as BiP (Grp78) or protein disulfide isomerase (PDI) remain largely unaltered (15–17).

To gain further insights into the nature of these peculiar ER structures, we set out to develop a more comprehensive understanding of their protein composition by comparing the localization of selected ER markers at 96 hpi during infection with either TB_148^HA^ (1), a recombinant HCMV strain TB40/E virus that expresses UL148 fused at its C-terminus to the nonapeptide epitope tag YPYDVPDYA from influenza A hemagglutinin (HA), or TB_159^HA^, a *UL148*-null comparator in which the UL148 homolog from rhesus cytomegalovirus, *Rh159*, likewise fused to a C-terminal HA tag, is expressed in lieu of UL148.

We considered the TB_159^HA^ virus to be an appropriate control for the following reasons. Firstly, Rh159 and UL148 exhibit ∼30% identity at the amino acid level and both glycoproteins localize to the ER and block cell surface presentation immune cell activating ligands; UL148 retains CD58, a ligand for CD2, while Rh159 retains NKG2D ligands of the MIC- and ULBP families (1, 9, 18). Secondly, Rh159 is expressed from TB_159^HA^ at comparable levels and with similar kinetics observed for UL148 from TB_148^HA^, and the two viruses replicate indistinguishably in fibroblasts (**FIG 2**). Moreover, as was the case in ectopic expression (10), Rh159 does not appear to activate the UPR to the same extent that UL148 does, as evidenced by lower levels of phospho-eIF2*α* and ATF4 during TB_159^HA^ infection (**FIG 2A**).

**FIG 2.**
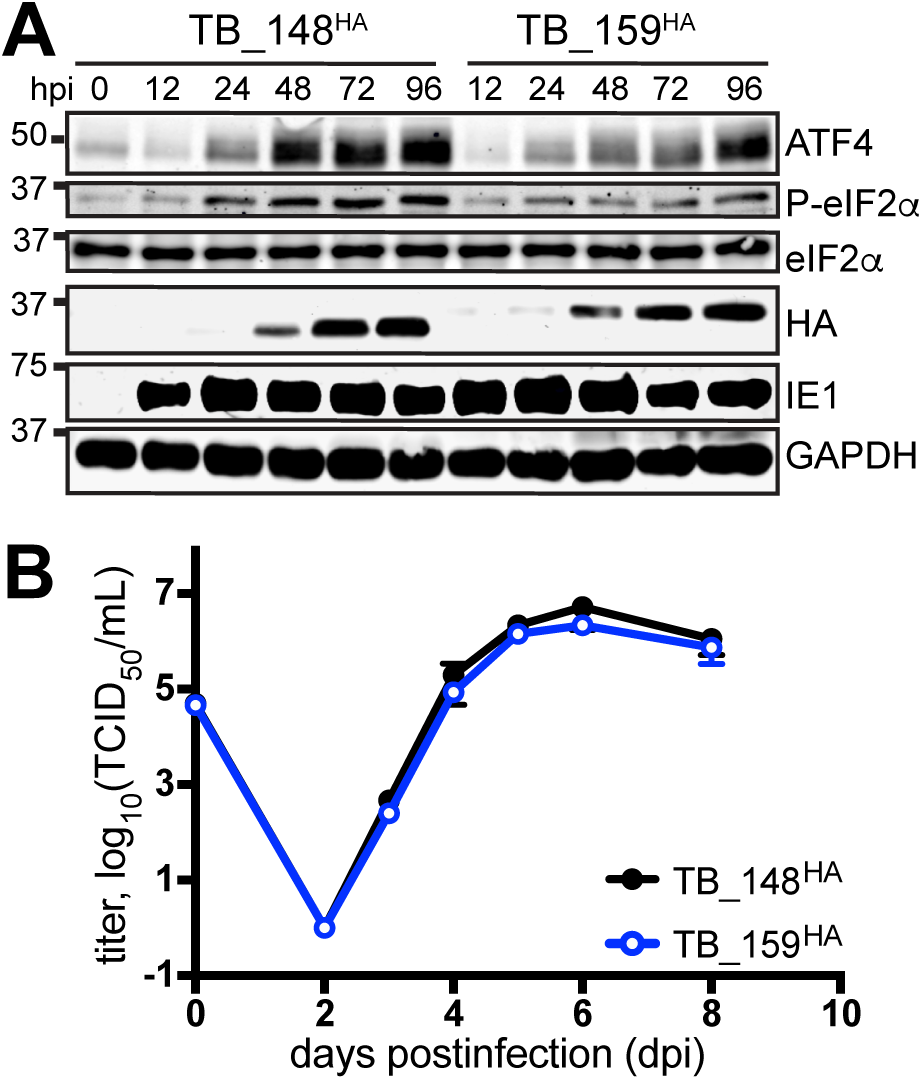
Characterization of TB_148^HA^ and TB_159^HA^ viruses. **(A)** Human fibroblasts were infected at MOI 1 with the indicated recombinant HCMVs. Cell lysate samples harvested at the indicated times postinfection (h postinfection, hpi) were analyzed by Western blot to detect HA-tagged UL148 or Rh159, the 72 kD viral nuclear antigen IE1-72 (IE1), and GAPDH. **(B)** Single-cycle viral replication kinetic curves from MOI 1 infected fibroblasts were plotted by determining the titer in tissue culture infectious dose 50 (TCID_50_) from supernatants collected at the indicated times postinfection.

ERQC markers failed to coalesce into unusual structures during infection with TB_159^HA^ (**FIG 3**). The structures likewise failed to occur in rhesus fibroblasts during infection with a recombinant rhesus cytomegalovirus (RhCMV) that we engineered to express an HA-tag fused at the C-terminus of Rh159 (**FIG S1**). The latter argues against the possibility that Rh159 somehow requires the context of rhesus cells to redistribute ER markers in a manner analogous to what is seen for UL148 during HCMV infection. Taken together, the data suggest that UL148 and Rh159 authentically differ in their effects on the secretory pathway.

**FIG 3.**
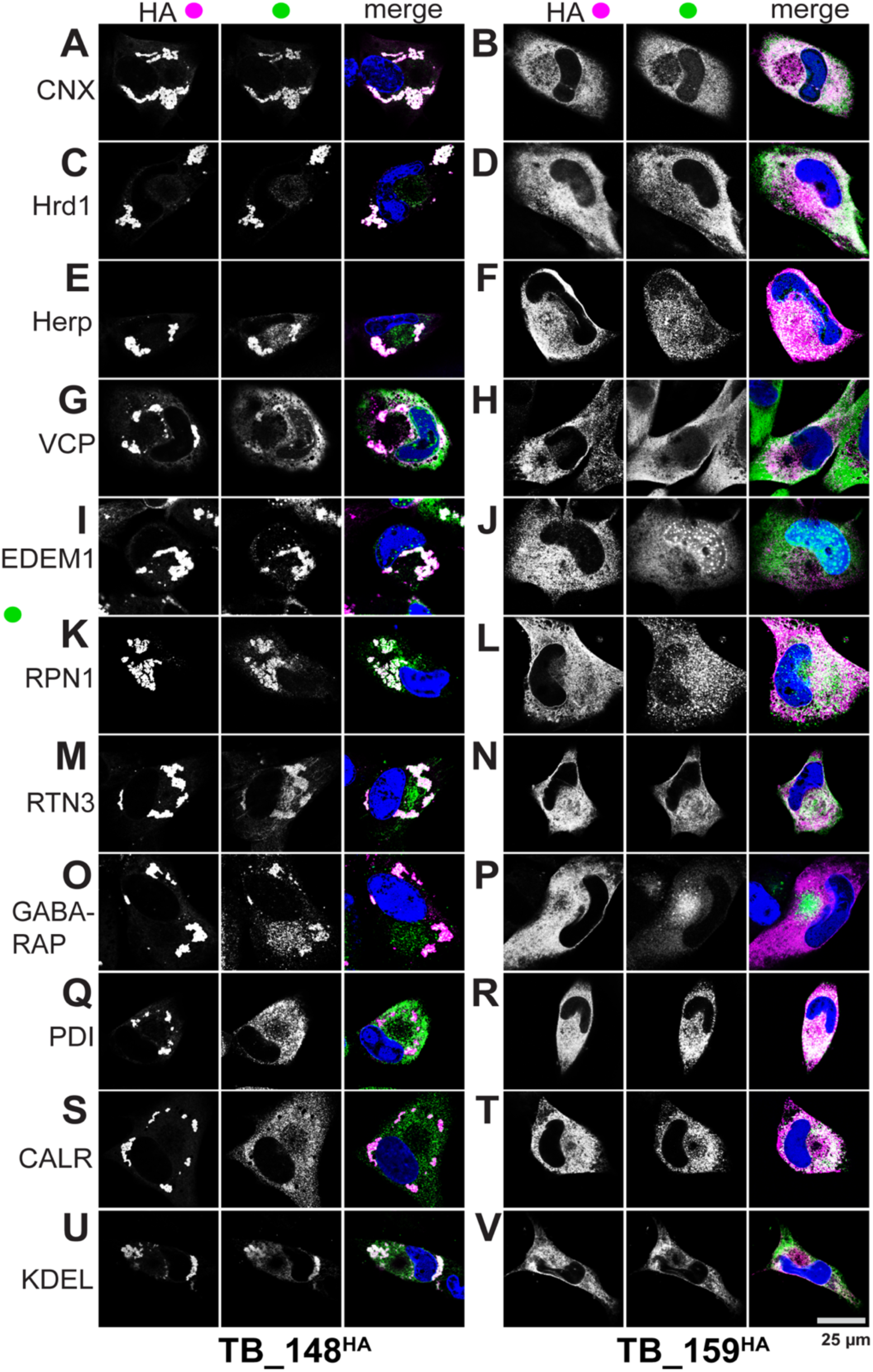
UL148 localizes to unusual ER compartments that are enriched for glycoprotein quality control markers. Fibroblasts infected at MOI 1 with either TB_148^HA^ (panels: **A, C, E, G, I, K, M, O, Q, S, U**) or TB_ 159^HA^ (panels: **B, D, F, H, J, L, N, P, R, T, V**) were fixed at 96 h postinfection, and imaged by confocal microscopy after co-staining with antibodies specific for HA (UL148 / Rh159, magenta in merge) and the indicated cellular markers (green in merge). DAPI (blue) counterstaining is shown in merged image.

In cells infected with TB_148^HA^, HA antibody staining indicated localization of UL148 to globular structures, as expected (1). Antibody signals from indirect confocal immunofluorescence detection of cellular ER resident proteins involved in ERQC, including calnexin, Hrd1, SEL1L, Herp, valosin containing protein (VCP, p97), and EDEM1, strongly co-localized with signals from HA-tagged UL148 (**FIG 3-4**). These results indicate that a number of ERQC factors coalesce with UL148 to form prominent globular structures in infected cells. In contrast, we observed uniform ER staining patterns for PDI and calreticulin (CALR), indicating that these ER markers do not localize to the UL148 structures (**FIG 3Q, 3S**). Intriguingly, antibody signals detecting reticulon 3 and ribophorin 1, which are markers for smooth ER and rough ER, respectively, each appreciably co-localized with UL148 (HA) signal at the induced structures (**FIG 3K, 3M**), which indicates that the structures may involve both rough and smooth ER. LC3B failed to co-localize with the structures (**FIG S2**), as might be expected given that the virus inhibits macroautophagy at late times during infection (19, 20). Nevertheless, our results suggested that the induced structures are enriched for a related mammalian ATG8 ortholog, GABARAP (**FIG 3O**).

**FIG 4.**
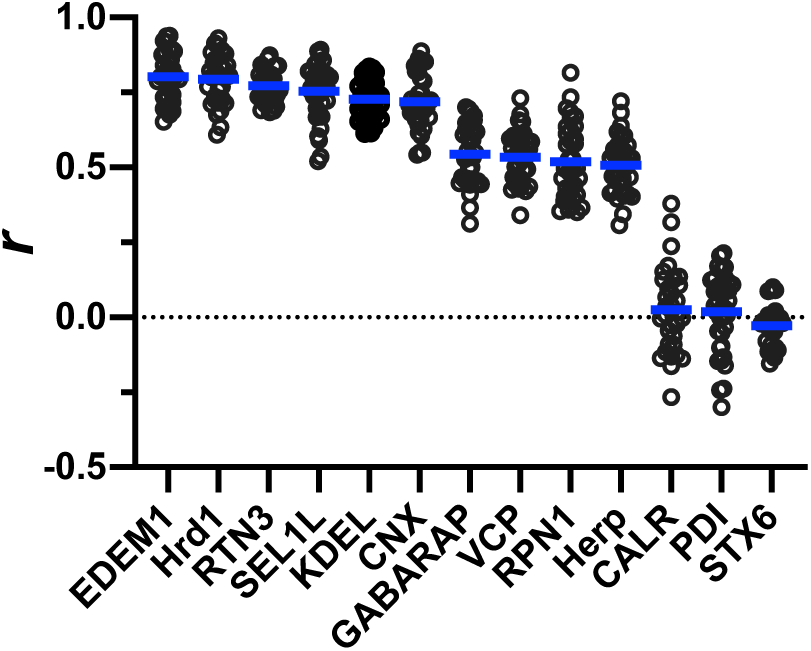
Quantification of co-localization between UL148 and cellular markers. A Pearson’s correlation coefficient (*r*) was calculated using NIH ImageJ software to estimate the degree of co-localization between UL148 (HA signal) and each of the indicated cellular markers. A minimum of 30 cells were analyzed per marker. The arithmetic mean for each co-localization analysis result is shown as a blue line, and data points for individual cells analyzed are plotted as circles.

In cells infected with TB_159^HA^, all of the ER markers we examined showed uniform, reticular staining, as did the anti-HA signal detecting Rh159 (**FIG 3B, D, F, H, J, L, N, R, T, V**). EDEM1, in addition to showing reticular staining, also labeled puncta associated with the nucleus (**FIG 3J, 3I**), which may represent enriched levels of the protein at the rough ER membranes associated with the nuclear envelope, although we cannot exclude the possibility of spurious intranuclear staining. GABARAP antibodies stained the cVAC in TB_159^HA^ infected cells, but also showed a much weaker diffuse signal throughout the cytosol. Even though we took steps to block viral Fc receptors, which can cause rabbit antibodies to non-specifically label the cVAC (21), it is plausible that GABARAP antibody signal from the cVAC reflects incomplete blocking of viral Fc receptors. For both viruses, antibodies specific for the TGN marker syntaxin-6 (STX6) and gH, as expected, stained the juxtanuclear cVAC structure (12, 13, 22), a post-ER compartment that excludes UL148 (1) (**FIG S2**).

To quantify the degree of co-localization with UL148, we calculated Pearson’s correlation coefficients from a minimum of thirty TB_148^HA^ infected cells per staining condition (fixation at 4 dpi), comparing the signal overlap for each ER marker to the HA signal from UL148. The correlation coefficients (*r*) for staining patterns from antibodies specific for EDEM1, Hrd1, reticulon-3 (RTN3), SEL1L, the ER retention motif KDEL, and calnexin ranged from 0.8 to 0.72. These results suggest that these proteins, and lumenal ER resident proteins such as BiP (GRP78) and GRP94 that contain a C-terminal KDEL motif (EKDEL), extensively co-localize with UL148 at the induced structures (**FIG 4**). Meanwhile, GABARAP, VCP, ribophorin-1 (RPN1), and Herp showed *r* values in the range of 0.54 to 0.51, indicating a moderate degree of co-localization with UL148. However, CALR and PDI, which like BiP, are lumenal ER residents, and the TGN marker syntaxin-6, gave *r* values of close to zero, indicating that these markers do not appreciably co-localize with UL148, as is consistent with our immunofluorescence data suggesting that these markers negligibly associate with the unusual structures. From these results, we conclude that the UL148-dependent ER structures are enriched for cellular markers involved in glycoprotein quality control. In this regard, the structures resemble the “ER quality control (ERQC) compartments” described by G. Lederkremer and colleagues (15, 16, 23). We further conclude that Rh159 cannot substitute for UL148 to redistribute ERQC markers.

### VCP and Hrd1 are recruited to incipient UL148 ER structures prior to calnexin

To determine whether there might be differences in the kinetics of recruitment of ERQC markers during formation of the ER structures, we examined a series of time points from one to four days post-infection (dpi) with wildtype HCMV strain TB40/E carrying HA-tagged UL148, staining for three different ERQC markers, calnexin, Hrd1, and VCP, alongside UL148 (HA). At 1 dpi the signal from UL148 was only faintly detected, as expected (1), while each of the ER markers was readily detected, providing a readout of their staining patterns prior to being substantially perturbed by UL148 (**FIG S3**). At 2 dpi, we detected robust anti-HA signal, indicating the presence of UL148. At this time point, UL148 exhibited intense signal at small globular puncta, which we interpret to represent incipient UL148 structures, as well as more diffuse staining of a reticular structure consistent with undisturbed ER. Calnexin did not appreciably co-localize with the UL148 puncta until at least 3 dpi, and the structures were not readily visualized by calnexin staining until 4 dpi (**FIG S3**). In contrast, signals from Hrd1 and VCP staining were sufficient to mark the UL148 puncta by 2 dpi, which suggests that these markers may co-localize with UL148 at earlier points during the genesis of the ER structures. Notably, the appearance of Hrd1 at the structures prior to calnexin is consistent with our previous results showing that UL148 co-purifies from infected cells with SEL1L, an adaptor subunit for Hrd1 (8). These results imply that basal elements of the ERAD machinery, exemplified by Hrd1 and VCP, might be recruited to UL148-induced ER structures prior to calnexin.

### Visualization of UL148-induced ER structures by electron microscopy

To discern the ultrastructural appearance of the UL148 structures and to formally ascertain their relationship to the ER, we carried out transmission electron microscopy (TEM) imaging of wildtype (TB_WT) and *UL148*-null (TB_148_STOP_) infected fibroblasts at 5 dpi. In high pressure frozen and freeze-substituted infected fibroblasts, cells with a high density of viral nucleocapsids within the nucleus were selected for analysis, as this feature indicates late time points during infection when UL148 is abundantly expressed. The TEM results revealed prominent globular and oblong structures in the cytoplasm of wildtype virus infected fibroblasts, but not *UL148*-null infected controls (**FIG 5-6**). The structures stand out for their high electron-density, which may reflect the abundance of ERQC proteins together with UL148 in these structures. Under higher magnification, these areas are characterized by accumulations of densely packed ruffled membranes and membranous material, which appear to either be collapsed ER, or ER tubules in very close association with each other. The membrane accumulations were associated with smooth and partially rough ER structures of seemingly enlarged cisternal space. Further, tomographic reconstruction of scanning transmission electron microscopy (STEM) data suggest that densely packed ER cisternae within the structures are interconnected and continuous in three-dimensional space (**FIG 7B-C, Movie S1**).

**FIG 5.**
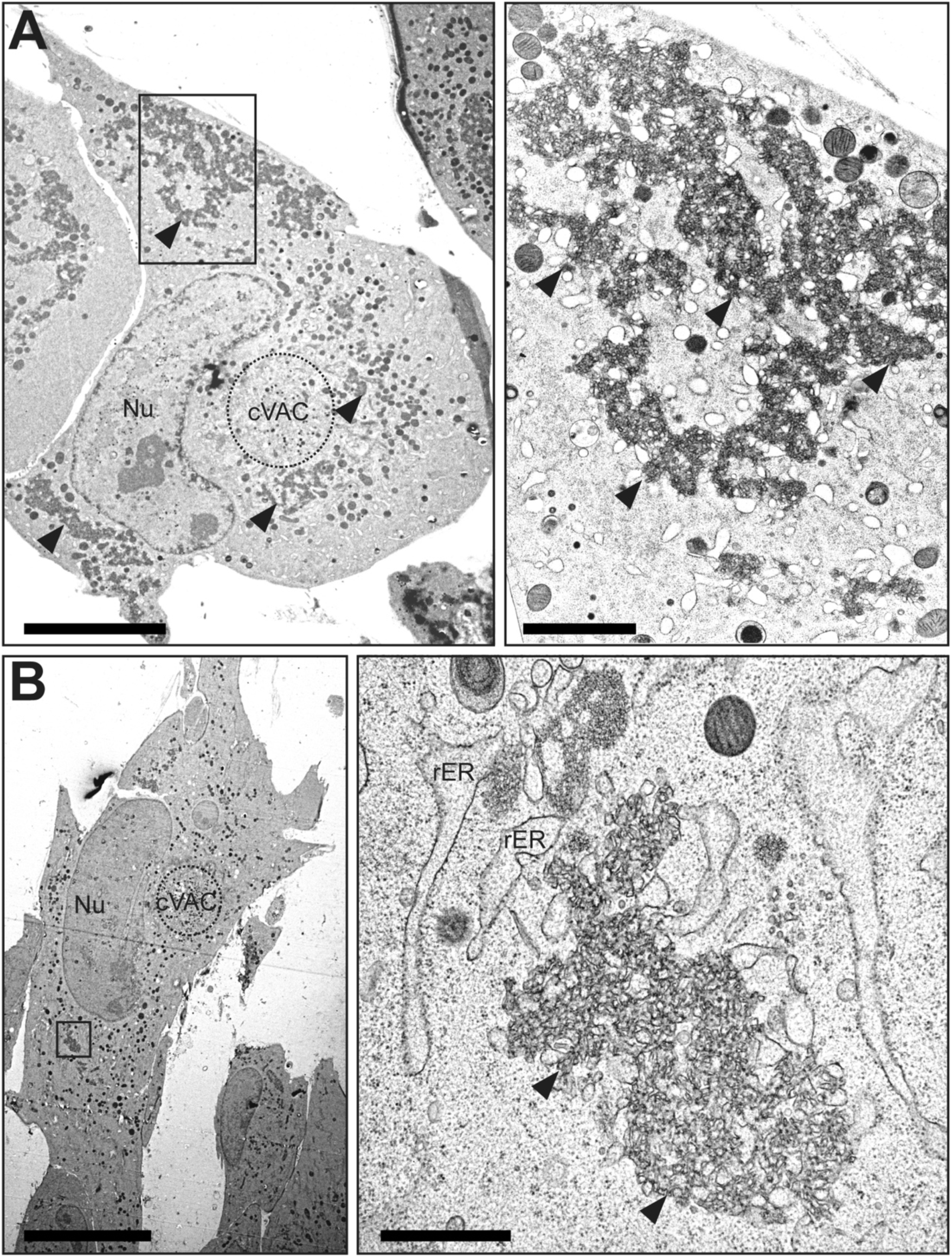
TEM of ER structures in wild-type HCMV infected cells. Human fibroblasts infected with wildtype HCMV (TB_WT) were fixed by high-pressure freezing and freeze substitution at day 5 postinfection and imaged using TEM. Panels (**A**) and (**B**) show cell overview at left. For each cell, the boxed region is shown at higher magnification. Scale bars; left panels: 10 µm, right panels: 2 µm. rER: rough ER; Nu: nucleus; cVAC: cytoplasmic viral assembly compartment. Solid arrowheads indicate the UL148-dependent ER structures of interest.

**FIG 6.**
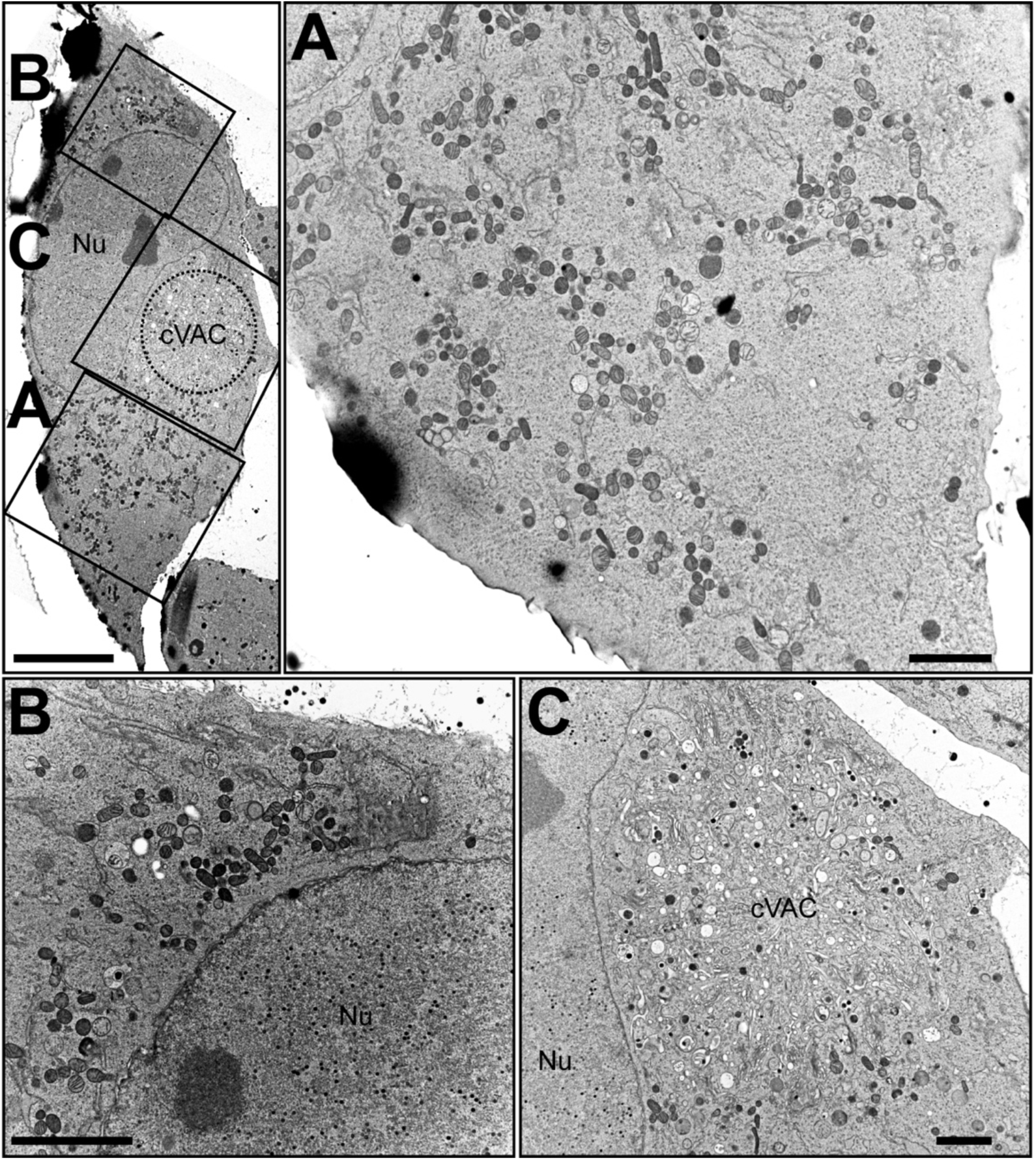
TEM of *UL148*-null HCMV infected cells. Human fibroblasts infected with a *UL148*-null mutant (TB_148_STOP_) were fixed by high-pressure freezing and freeze substitution at day 5 postinfection and imaged using TEM. At the upper left an overview panel of a representative cell is shown with panels (**A)**, **(B)**, and **(C)** each boxed. The boxed regions are expanded at higher magnification at right (**A**) and below (**B, C**). Scale bars for upper left overview panel: 10 µm, for zoomed panels: 2 µm. rER: rough ER; Nu: nucleus; cVAC: cytoplasmic viral assembly compartment.

**FIG 7.**
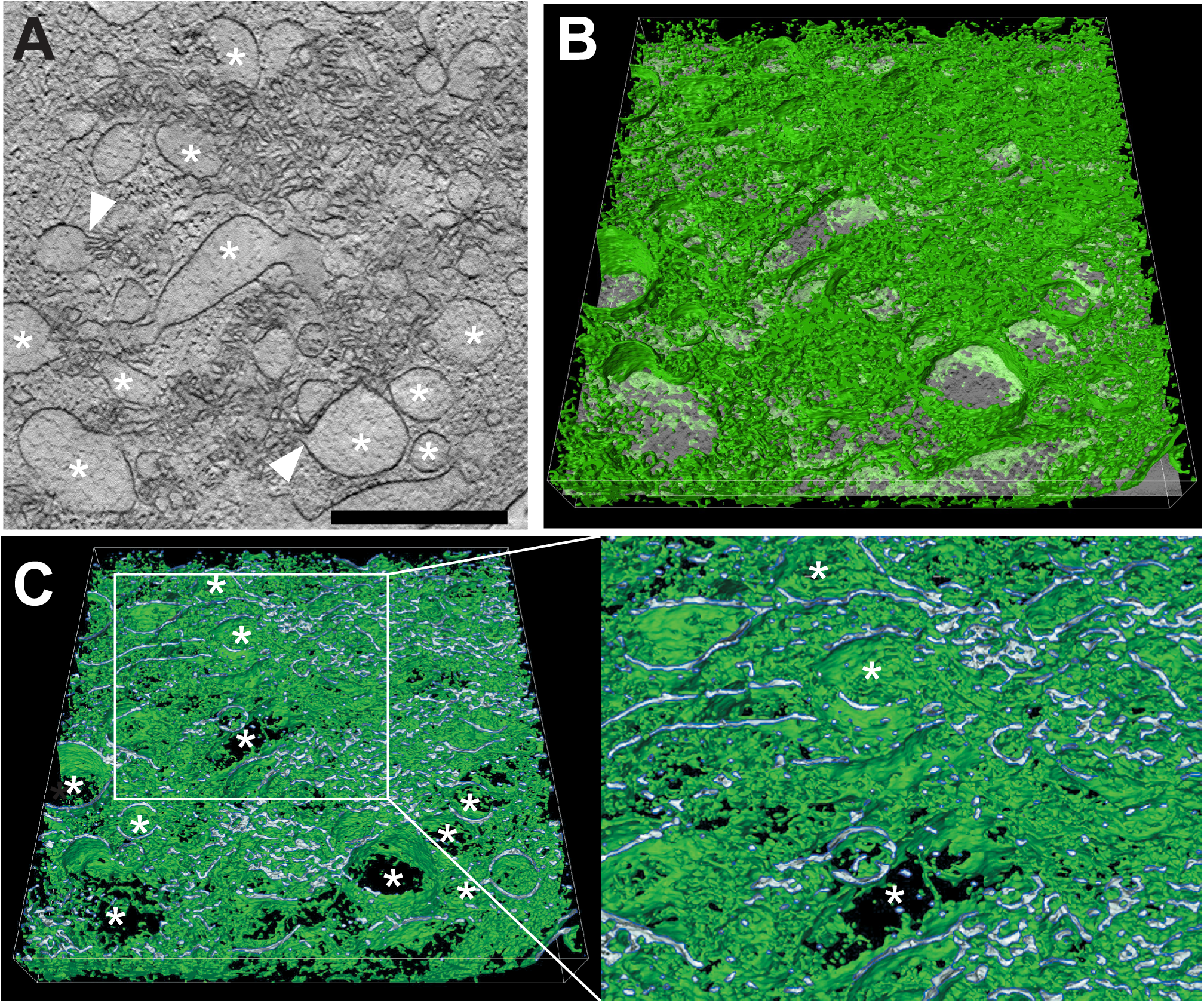
STEM tomography of UL148-dependent ER structures in HCMV infected cells. Human fibroblasts infected with wildtype HCMV (TB_WT) were fixed by high-pressure freezing and freeze substitution at day 5 postinfection and tomograms were recorded by STEM. (**A**) Shows a virtual section through the tomogram of a virus-induced membranous structure; asterisks denote ER structures of distended lumenal space. White arrowheads indicate sites at which membranes originating from distended ER cisternae continue into areas of involuted collapsed ER. Scale bar: 1 µm. (**B**) Shows the same virtual section as in (A) tilted and with a 3D visualization of the membranous network (green) of the entire tomogram. The same distended ER cisternae as in (A) are marked by asterisks. (**C**) Shows a cross section through (B) to visualize the membrane profile of the membranous structures. The region delimited by a white box is shown in a higher magnification on the right. Finer detail of the enlarged ER cisternae and the connections between them are readily visible; asterisks indicate the same distended ER cisternae as in panels (A) and (B). Also, see SI Movie S1. Scale bar: 1 µm.

### UL148 accumulates in a detergent-insoluble form during infection

Disease-associated variants of certain cellular proteins, such as the A103E mutant of the calcium channel Orai1, and the E342K “Z” variant of alpha-1 antitrypsin (Z A1AT), localize to anomalous ER structures reminiscent of those we observe to depend on UL148 (17, 24–27). A103E Orai1 and Z A1AT are found to accumulate in detergent-insoluble forms within the ER or within ER membranes, respectively, which indicates aggregation or polymerization of these proteins, and suggests a mechanism for the formation of ER structures by the proteins (17, 24, 27, 28). We therefore set out to determine whether differences in solubility might correlate with the differing potentials UL148 and Rh159 to cause ER reorganization and to activate the UPR (10). To address this question, we infected fibroblasts with TB_148^HA^ or TB_159^HA^ at MOI 1 and at various times post-infection prepared cell lysates in radioimmunoprecipitation assay (RIPA) buffer. After subjecting the lysates to high-speed centrifugation, we examined the relative levels of UL148 and Rh159 in the detergent-soluble supernatant and detergent-insoluble pellet fractions.

A substantial portion of UL148 was detected from the detergent-insoluble fractions at all time points tested (**FIG 8**). Furthermore, the percentage of UL148 detected within the insoluble fraction increased over time. In contrast, Rh159 was found only in the detergent-soluble fraction. From these results, we conclude that UL148 accumulates in a detergent-insoluble form during infection, and that Rh159 does not do so. Because the anti-HA immunoreactive band detected in both the soluble and insoluble fractions from TB_148^HA^ infected cells showed a relative mobility of ∼35 kD, which matched that expected for the mature, endoH-sensitive glycoprotein (1), these findings argue that UL148 may form aggregates or polymers within the ER.

**FIG 8.**
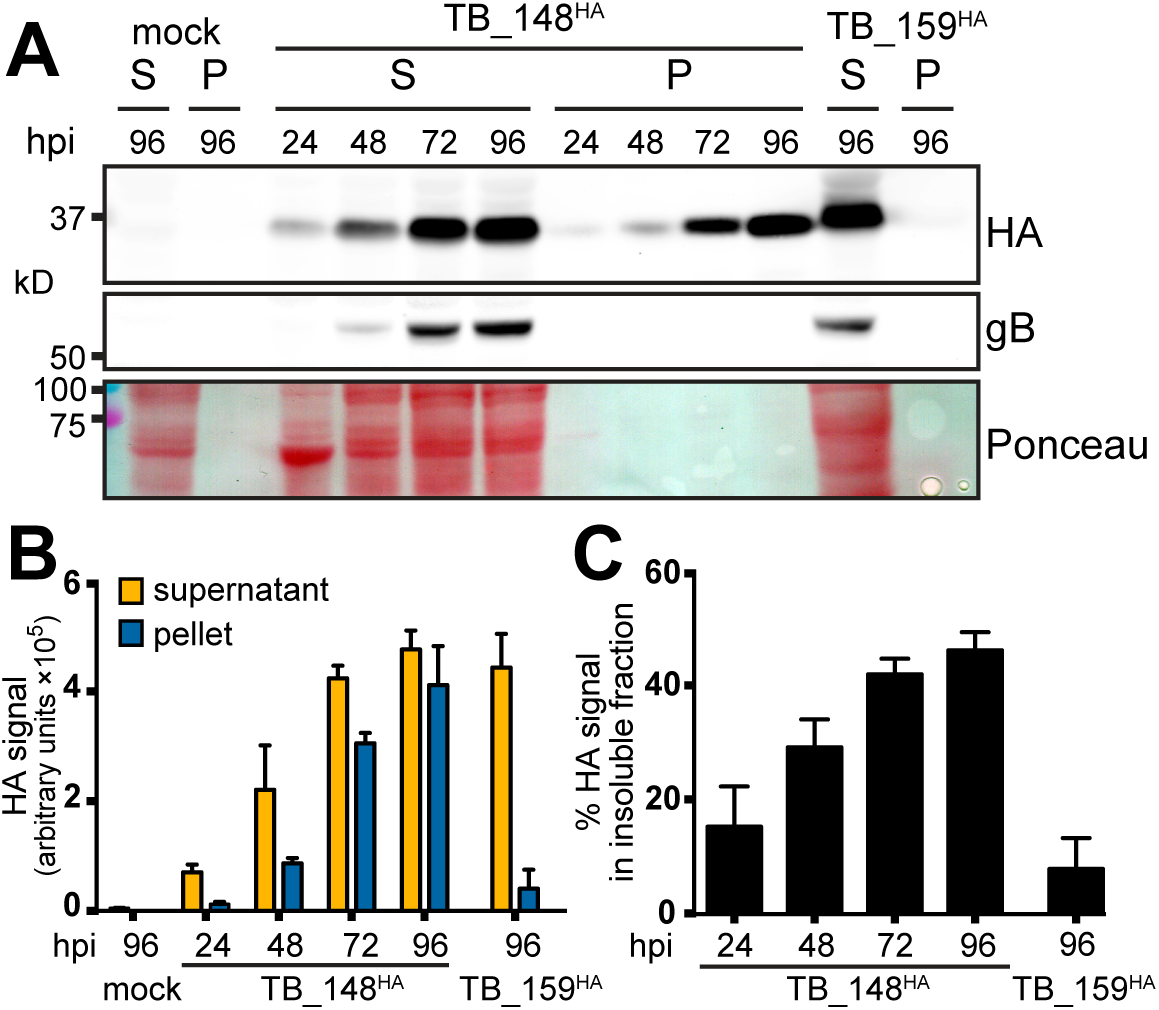
Solubility analysis of UL148 and Rh159. (**A**) Human fibroblasts were infected at MOI 1 TCID_50_/cell with HCMV strain TB40/E derived viruses TB_148^HA^, which expresses UL148 fused at its C-terminus to an HA-epitope tag, or TB_159^HA^, which lacks *UL148* and instead expresses rhesus CMV Rh159 carrying a C-terminal HA tag. At the indicated times postinfection (hpi), infected cells were collected in radioimmunoprecipitation (RIPA) lysis buffer, centrifuged at 21,000 *× g* for 30 min, after which supernatant (S) and pellet (P) fractions were boiled in gel loading buffer containing 2% sodium dodecyl sulfate (SDS). Equivalent portions of supernatant and pellet were resolved by SDS-polyacrylamide gel electrophoresis, transferred to a nitrocellulose membrane for detection of protein species immunoreactive to antibodies against HA epitope (HA) and HCMV glycoprotein B (gB) and for total protein signal using Ponceau S reagent (Ponceau). (**B-C**). Signal intensity of fluorophore-conjugated secondary antibodies in anti-HA Western blots were measured from three independent biological replicates of the experiment shown in (A); error bars indicate standard deviation. (**B**) The fluorescent signal for each infection time point condition (y-axis is in arbitrary units, in hundred thousands). (**C**) The amount of signal found in the insoluble (pellet) fraction relative to the total signal (pellet plus supernatant) for each infection time point are plotted as percentage values.

### UL148 is sufficient to compartmentalize the ER

Our data thus far demonstrate that UL148 is necessary during infection to cause redistribution of cellular ER markers for glycoprotein quality control processes, such that substantial portion of the ERQC machinery appears to become sequestered away from the rest of the organelle into novel membranous structures. Because a number of viral proteins that remodel the ER during infection are sufficient to alter ER morphology when ectopically expressed [e.g. (29),(30), reviewed in (31)], we wondered whether UL148 expression would be sufficient to induce ER structures akin to those seen during HCMV infection. Making use of “tet-on” ARPE-19 epithelial cells that inducibly express either UL148 or Rh159, each carrying a C-terminal HA-tag (32), we induced transgene expression for 48 h and subsequently stained for various cellular markers for the ER, for ERQC as well as for the ATG8 family proteins LC3B and GABARAP, which play roles in macroautophagy and related processes.

In cells expressing UL148 (i148^HA^), we observed that calnexin, Hrd1, EDEM1, and VCP co-localized with UL148 at prominent globular structures reminiscent of those observed during infection (**FIG S4**). The respective rough and smooth ER markers ribophorin 1 (RPN1) and reticulon 3 (RTN3), as well as the ATG8 proteins LC3B and GABARAP likewise co-localized to the UL148-induced structures. However, PDI and CALR each failed to substantially co-localize with UL148 at the structures (**FIG S4**), or at best showed only limited co-localization, consistent with our results from infected cells (**FIG 3-4**). Signals from antibodies specific for the KDEL (EKDEL) motif important for ER retrieval of luminal ER residents, such as BiP and GRP94, likewise showed only limited co-localization with UL148. The PDI and KDEL results suggest that large regions of the ER are not involved in the UL148 structures.

In cells expressing the UL148 homolog Rh159 (i159^HA^), we detected uniform cytoplasmic distribution of ER markers, as well as of HA-tagged Rh159, similar to what we observed during infection with the recombinant HCMV TB_159^HA^ (**FIG 3**). Furthermore, in this setting the staining patterns for the ATG8 family proteins LC3B and GABARAP failed to indicate any notable structures. Consistent with our previous study showing that UL148, but not Rh159, is sufficient to activate the UPR (10), we observed accumulation of ATF4 and phosphorylated eIF2*α* following doxycycline (dox) induced expression of UL148 but not Rh159 (**FIG 9A**). Importantly, the intensity of HA signals detecting UL148 and Rh159 indicated that both ER resident immunevasins accumulate at roughly comparable levels following dox induction. This argues against the possibility that gross protein expression differences accounts for the differing effects of UL148 and Rh159 on UPR activation and on the staining patterns for ER markers.

**FIG 9.**
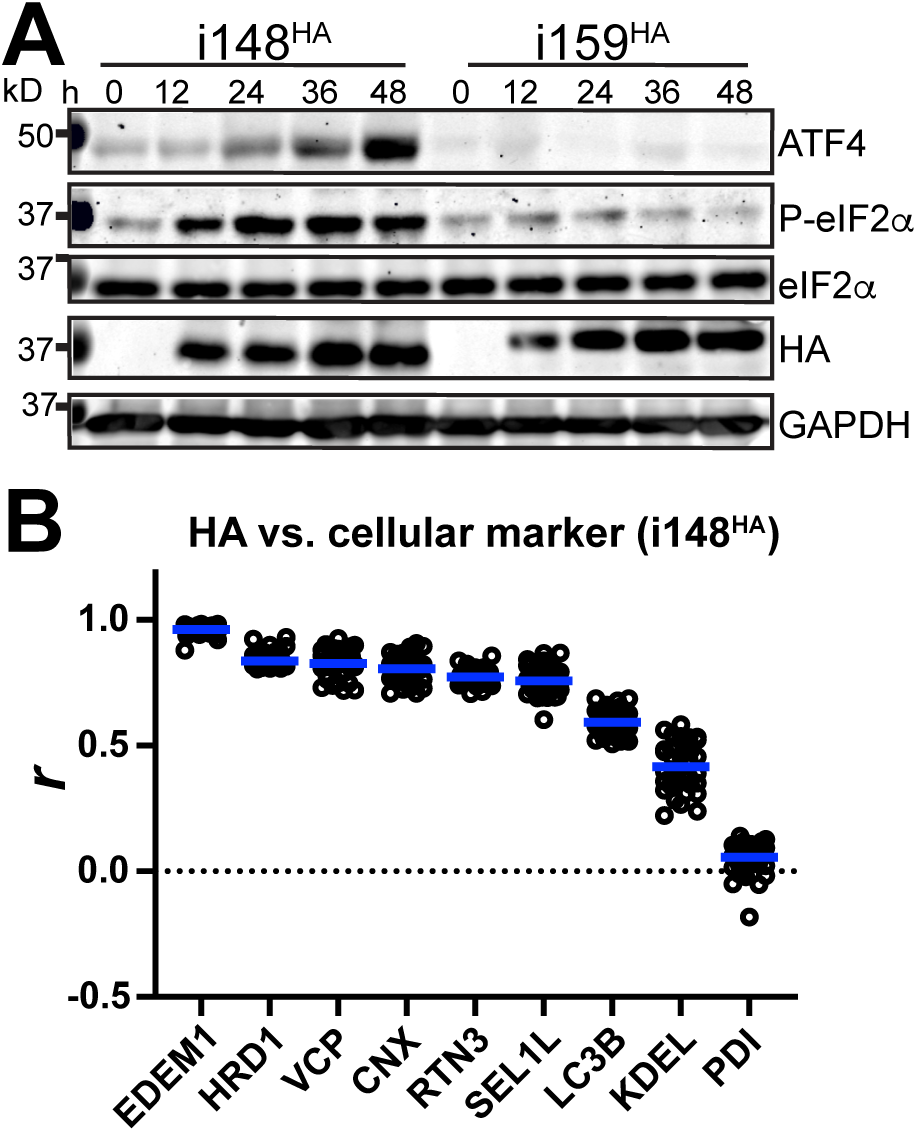
ISR activation accompanies redistribution of ER markers during ectopic expression of UL148. **(A)** Lysates of ‘tet-on’ ARPE-19 cells expressing either UL148 or Rh159 fused to an HA tag, i148^HA^ and i159^HA^, respectively, were collected at the indicated times post doxycycline (dox) induction and analyzed by Western blot for the expression of the indicated proteins and for the abundance of eIF2*α* phosphorylated at Ser51 (P-eIF2*α*) using a phospho-specific antibody. (**B**) Pearson’s correlation coefficient (*r*) values were calculated using NIH ImageJ software estimate the degree of co-localization between UL148 (HA signal) and the indicated cellular markers. A minimum of 30 cells were analyzed per marker. Arithmetic means for each co-localization analysis result are shown as blue lines, and data points for individual cells analyzed are plotted as circles.

Pearson’s correlation coefficient values were calculated to quantify the extent of colocalization between UL148 and various cellular markers: EDEM1, Hrd1, VCP, calnexin (CNX), RTN3, SEL1L, LC3B, and PDI (**FIG 9B**). The results quantitatively buttress the confocal immunofluorescence results from these cells (**FIG S4**). We therefore conclude that no other HCMV gene products are required for UL148 to cause ERQC factors to segregate into discrete compartments. Hence, we conclude from these findings that UL148 is sufficient to cause large-scale reorganization of ERQC markers. Moreover, these results suggest that reorganization of ER markers into discrete structures may be related to the propensity of UL148 to induce ER stress.

To determine whether UL148-dependent ER remodeling could be visualized in real-time using live cell imaging, we constructed new “tet-on” lentiviral vectors that inducibly express either UL148 or Rh159 fused at their predicted C-terminal cytoplasmic tails to the enhanced green fluorescent protein (GFP) from *Aequorea victoria* (33). Following lentiviral transduction, we isolated puromycin-resistant ARPE-19 and subsequently subjected to fluorescence-activated cell sorting (FACS) to enrich for cells that expressed GFP signal following dox treatment. The resulting cell populations, which inducibly express either UL148-GFP or Rh159-GFP, were designated i148^GFP^ and i159^GFP^, respectively.

In live cell imaging studies, we observed that the GFP signal in dox-induced i148^GFP^ cells first appeared in a reticular, largely uniform pattern, which was readily visible at 5 h post-induction. However, by 9 h post-induction punctate signals began to appear. These puncta were observed to traffic to sites of large-scale accumulation, where large fluorescent structures progressively increased in size up until the termination of the experiment at 19 h (**FIG S5, Movie S2**). In contrast, i159^GFP^ cells exhibited a largely diffuse, reticular GFP signal at all time points monitored subsequent to transgene induction (**FIG S5, Movie S3**). The live cell imaging results thus recapitulated the differences in the HA-staining patterns observed between dox-induced i148^HA^ and i159^HA^ cells following fixation, as well as those between HCMV expressing HA-tagged UL148 versus *UL148*-null comparator viruses (**FIG 1, 3, Fig S1**).

### UL148-GFP structures exhibit poor recovery of fluorescence after photobleaching

To determine whether the protein contents of the UL148 induced structures exhibits decreased mobility compared to non-perturbed ER regions, we carried out fluorescence recovery after photobleaching (FRAP) studies. We photobleached regions of GFP signal in i148^GFP^ or i159^GFP^ ARPE-19 cells that had been induced for transgene expression for 24 h, and then monitored recovery of fluorescence over a 5 min time period. In Rh159-GFP expressing cells (i159^GFP^), fluorescence nearly recovered to pre-bleach levels within 3 min, and by 5 min fully recovered (**FIG S6**). In contrast, when we photobleached a prominent UL148-GFP structure, the GFP signal failed to appreciably recover fluorescence during the same 5 min time period. Nonetheless, regions of reticular GFP signal from an i148^GFP^ cell, which presumably represent ER regions not involved in an anomalous structure, recovered fluorescent signal with kinetics similar to those observed during Rh159-GFP expression. These results indicate that bleached UL148-GFP within the structures cannot rapidly be exchanged with UL148-GFP from other portions of the organelle, which suggests that the induced UL148 structures do not exchange their contents as efficiently as unperturbed ER.

### Ectopically-expressed UL148 is degraded by proteasomes and not by autophagy

The UL148 ER structures were found to be enriched with proteins such as Hrd1, SEL1L, EDEM, and VCP (**FIG 3-4, 9B, FIG S4**), which are posited to play key roles in ERAD, a process during which misfolded glycoproteins are recognized, processed, and dislocated across the ER membrane for degradation at cytosolic proteasomes. However, we also detected elements of the machinery for autophagy in close association with the UL148 structures. Namely, the mammalian ATG8 ortholog GABARAP co-localized with UL148 during infection (**FIG 3-4**), and both GABARAP and another ATG8 ortholog, LC3B, co-localized with UL148 at the ER structures during ectopic expression of UL148 (**FIG 9B, FIG S4**). Therefore, we wished to determine whether UL148 is degraded by the proteasome, as would be consistent with conventional ERAD, or by the lysosome, which would suggest a role for autophagy-related processes, such as selective autophagy of the ER (34), in dispensing with UL148, and presumably, in resolving the ER perturbations.

As a first step, we conducted a live-cell imaging “washout” experiment in which i148^GFP^ cells were induced for transgene expression for 24 h, after which the growth medium containing the transgene inducing agent (dox) was replaced with medium lacking dox. During a 22 h imaging period following dox-washout, we observed the structures to become progressively smaller as the reticular and punctate GFP signals gradually abated (**SI FIG S7, Movie S7**). Because the results suggested that UL148-GFP structures could be resolved over time, we induced UL148-GFP in tet-on ARPE-19 cells (i148^gfp^) for 24 h, washed out the inducing agent (dox), and then applied either epoxomicin (20 µM), an irreversible inhibitor of the proteasome (35), or folimycin (115 nM), a proton pump inhibitor that blocks autophagosome maturation and impedes lysosome-dependent degradation (36, 37). Of the two inhibitors, only epoxomicin stabilized UL148 (**SI FIG S7**). Although folimycin treatment had no obvious effect on UL148-GFP levels, the drug markedly increased the levels of LC3B-II, as would be expected with the lysosomal proton pump inhibitor. From these results, we conclude that ectopically expressed UL148 is primarily degraded by proteasome-dependent pathway.

### UL148 requires the integrated stress response to cause ER reorganization

We previously reported that UL148 triggers the UPR during ectopic expression, and that UL148 contributes to activation of the PKR-like ER kinase (PERK) and inositol requiring enzyme 1 (IRE1) during infection (10). The literature suggests that formation of ERQC compartments requires PERK (38). PERK responds to ER stress by activating the integrated stress response (ISR). In particular, PERK phosphorylates eIF2*α* at Ser51, and the accumulation of phosphorylated eIF2*α* (eIF2*α*-P) globally attenuates mRNA translation whilst stimulating the translation of a subset of mRNAs, such as those encoding ATF4 and CHOP, which play roles in cellular adaptation to stress (39). Stress-regulated translation of such mRNAs involves small upstream open reading frames (uORFs) in their 5’ untranslated regions that ordinarily suppress translation under basal conditions (40, 41).

To examine whether ER remodeling in response to UL148 requires the ISR, we turned to a well characterized small molecule inhibitor of stress-regulated translation, ISRIB (42–46). ISRIB is thought to act as a “molecular staple” that holds the guanine nucleotide exchange factor eIF2B in an active decameric configuration (44–47), such that eIF2B will continue to generate ternary complex (eIF2•GTP•Met-tRNAi) necessary for new cycles of translational initiation, despite the presence of eIF2*α*-P. Because PERK is the kinase that phosphorylates eIF2*α* in response to ER stress (48, 49), we also tested for effects of the PERK inhibitor GSK2606414 (50). Having confirmed that ISRIB and GSK260641 do not negatively impact UL148 expression during dox-induction of UL148-GFP (**FIG 10**), we treated i148^GFP^ cells with 200 nM ISRIB, 1.1 µM GSK260641, or DMSO vehicle control, and carried out live cell imaging during induction of UL148-GFP expression.

**FIG 10.**
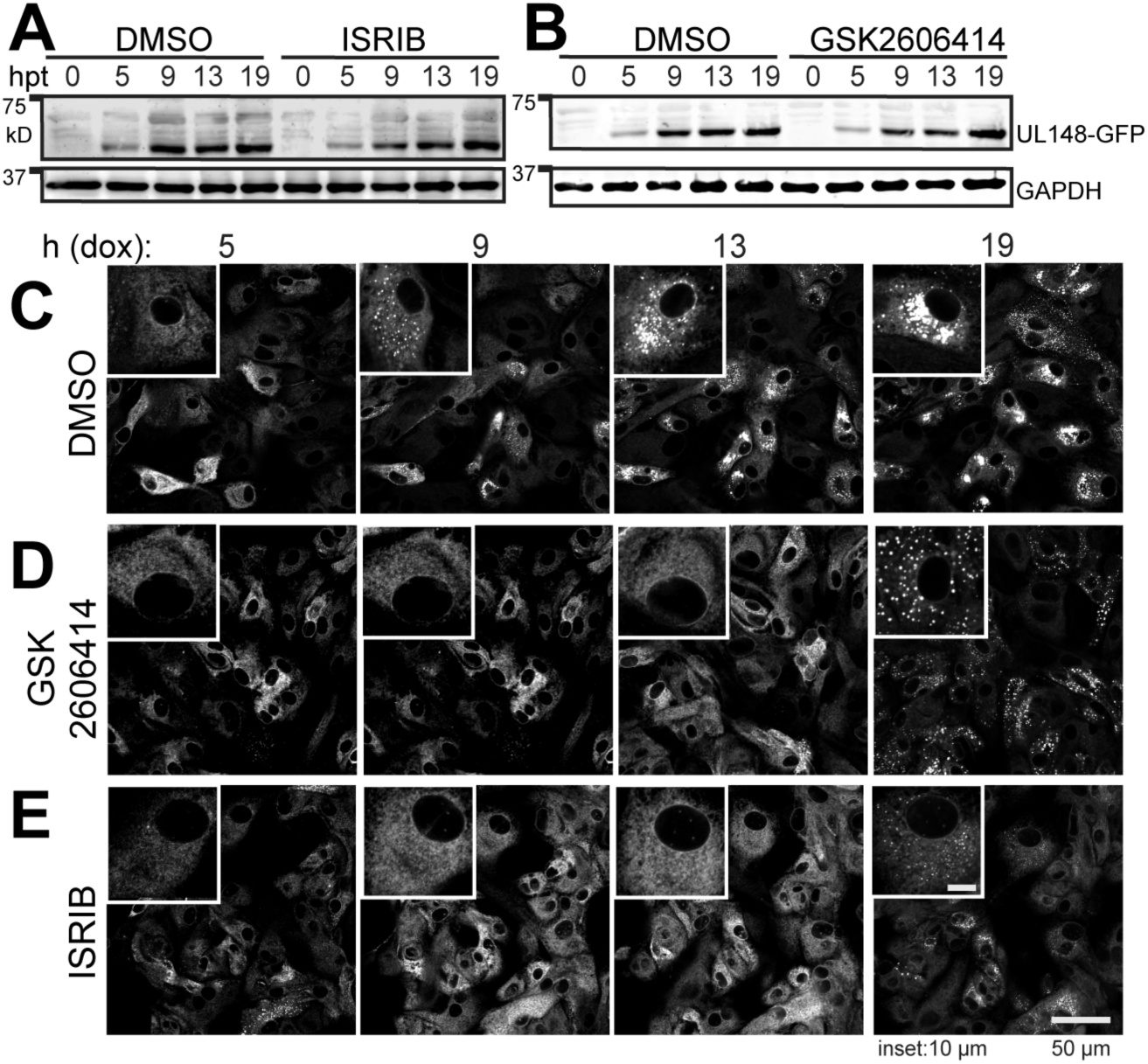
Inhibition of the integrated stress response impedes UL148-mediated ER remodeling. (A-B) “Tet-on” ARPE-19 epithelial cells that inducibly express UL148 fused to green florescent protein (gfp), i148^gfp^, were induced for transgene expression using 100 ng/mL doxycycline (dox) in the presence of DMSO carrier, ISRIB (200 nM), or GSK2606414 (1.1 µM) and monitored by anti-UL148 Western blot for expression of UL148-GFP over a series of time points (h post treatment with dox, hpt). (**C-E**) Live cell imaging of UL148-GFP expression patterns in the presence of ISRIB (200 nM), GSK2606414 (1.1 µM), or DMSO vehicle (0.01%). Main scale bar represents 50 µm. For inset panels at upper left of each image, which are magnified 2.4*×* relative to the main image, the scale bar represents 10 µm. Also see SI Movies S8-S10.

ISRIB and GSK6060414 virtually abolished the formation of UL148 puncta through 13 h post induction (dox addition), a time point at which the DMSO control condition showed abundant large structures (**FIG 10C-E, SI Movies S8-S10**). Based on our findings (**FIG 1, 3, 5, 7, 9B, S4**), we interpret the formation and subsequent large-scale aggregation of UL148-GFP puncta to faithfully indicate ER reorganization in response to UL148. Therefore, these results suggested to us that pharmacological inhibition of either the ISR or PERK prevent UL148 from remodeling the ER, which implies that ER remodeling in response to ER stress requires stress-regulated translation.

Since our results with ISRIB and GSK6060414 indicated that the ISR is required for ER remodeling to occur during ectopic expression of UL148, we next asked whether these pharmacological agents would prevent UL148-dependent ER remodeling in the physiologically authentic context of HCMV infection. Our live cell imaging studies from ISRIB and GSK6060414 treatments indicated that UL148-GFP puncta could begin to appear by ∼19 h post treatment (**FIG 10, SI Movies S9-S10**), which suggested that the pharmacological activity of the drugs might diminish over time. Thus to ensure that blockade of PERK and the ISR would be sustained over the 4 d period that it takes for structures to fully form during HCMV infection, we replenished drug treatments every 24 h, replacing the spent media with media containing freshly reconstituted ISRIB, GSK6060414, or DMSO vehicle control.

At 4 dpi, ER structures were readily visible in DMSO control treatment condition but virtually abolished in the ISRIB and GSK2606414 conditions (**FIG 11A**). To quantify these effects, we obtained Z-stacks from a minimum of 30 cells per treatment condition and used Imaris 3D image analysis software to calculate the percentage of Hrd1 signal that coalesced into discrete structures during infection. The results indicated highly significant differences between the DMSO carrier alone setting compared to treatments with either GSK2606414 or ISRIB (**FIG 11B**). In the DMSO control condition, on average, 10% of the Hrd1 signal was found to be associated with the UL148 structures (arithmetic mean; range 5.8% – 17.6%), as indicated by structures delimited by HA signal detecting UL148. In the presence of ISRIB or GSK2606414, however, these values were reduced to 2.8% (range: 0.69% – 7.1%) and 3.6% (range: 0.8%-6.5%), respectively. Reassuringly, roughly equivalent levels of UL148 were detected in Western blot analyses of protein extracts from drug treated and DMSO control conditions. Therefore, differences in UL148 expression are unlikely to explain the observed failure of UL148 and Hrd1 to coalesce into ER structures during inhibition of either the ISR or PERK. Meanwhile, ISRIB and GSK2606414 treatments both led to reduced levels of ATF4 accumulation, while the PERK inhibitor alone was able to reduce the levels eIF2*α*-P, as indicated by a phospho-specific antibody. Therefore, each of the drugs caused the expected effects on the PERK, eIF2*α*-P, and ATF4 axis (**FIG 11C-D**).

**FIG 11.**
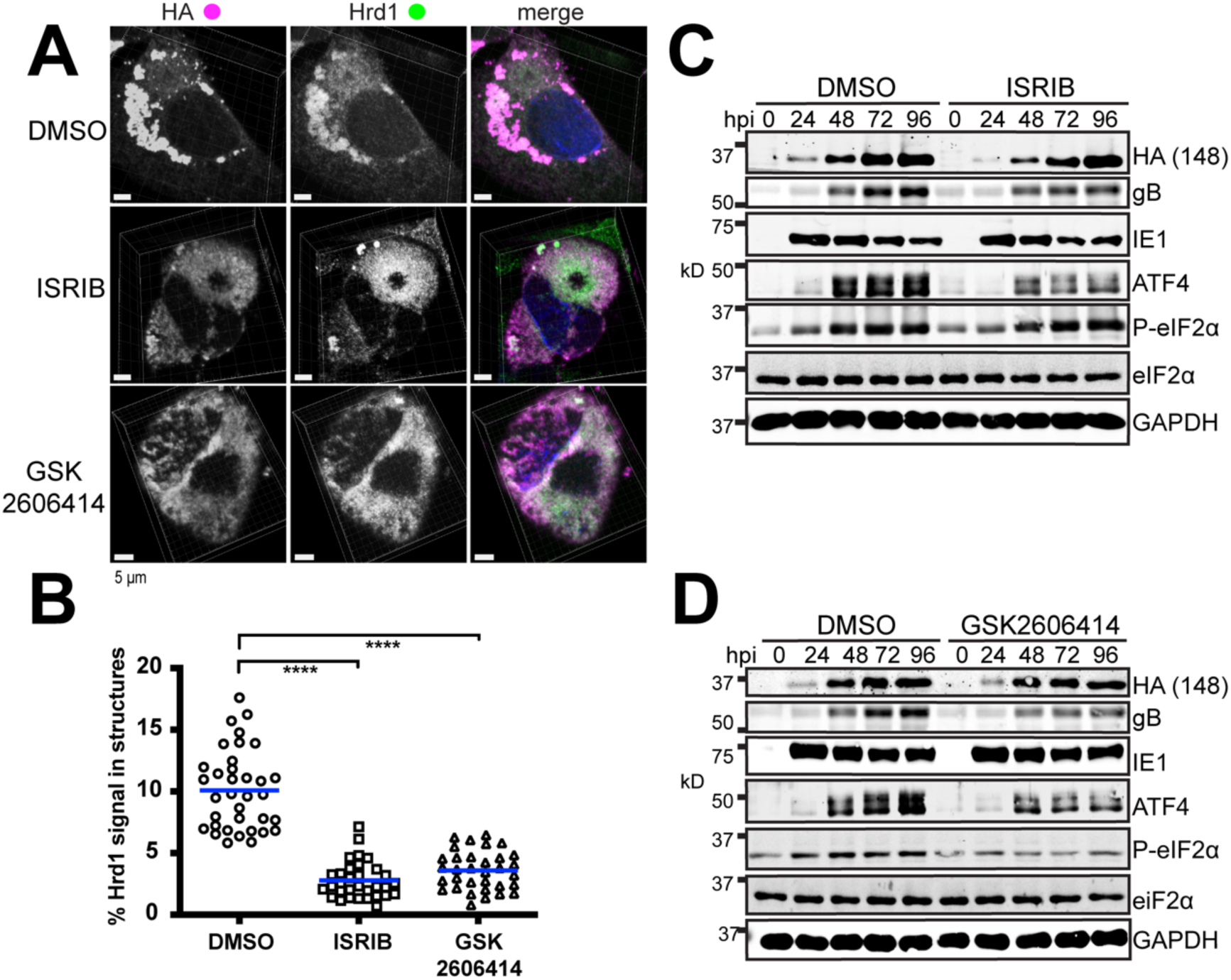
Inhibition of the ISR prevents the coalescence of Hrd1 and UL148 into discrete structures during infection. (**A**) Representative 3D maximum intensity projections of confocal imaging Z-stacks obtained from cells infected at MOI 1 for 96 h with HCMV strain TB40/E carrying HA-tagged *UL148* (TB_148^HA^) and maintained in the presence of ISRIB (200 nM), GSK2606414 (1.1 µM), or DMSO carrier alone (0.1% vol/vol). In merged images HA signal is shown in magenta, Hrd1 in green, and DAPI counterstaining in blue. (**B**) The percent of Hrd1 antibody signal involved in discrete structures at 96 hpi were calculated for a minimum of 30 cells per condition using Imaris x64 9.3.0 software. Statistical significance was determined using a one-way ANOVA followed by Tukey’s post-test; **** represents a P-value of <0.0001. The arithmetic mean for co-localization analysis results are shown as blue lines with data points for individual cells analyzed plotted as circles, squares or triangles, as indicated. (**C-D**) Western blot analyses of fibroblasts infected at MOI 1 with TB_148^HA^ and maintained in the presence of ISRIB (200 nM), GSK2606414 (1.1 µM), or DMSO carrier alone (0.01%); hpi: h post infection (hpi). Note: a phospho-specific antibody was used for detection of eIF2*α* phosphorylated at Ser51 (P-eIF2*α*).

A modest reduction in the levels of the viral envelope glycoprotein B (gB) was observed during treatments with ISRIB or GSK2606414. This may suggest that the ISR is required for optimal expression of the certain viral envelope glycoproteins. Even though UL148 promotes ATF4 expression, likely via PERK activation (10), viruses that lack *UL148* cause high levels of ATF4 accumulation at late times during infection, which is when viral late gene products, such a gB, are expressed at their highest levels (10). Hence, the activity of PERK and/or of the ISR may be required for optimal expression of viral envelope glycoproteins. Regardless, the PERK inhibitor and ISRIB each prevented the appearance of discrete UL148 ER structures while failing to substantially affect UL148 expression. Therefore, the effects of the two inhibitors on UL148-dependent ER reorganization were consistent with those seen during ectopic expression of the protein. Taken together, our findings argue that the ISR is required for UL148 to cause remodeling of the organelle.

## DISCUSSION

The ER structures that we have identified are noteworthy in several regards. Firstly, this example of virally-induced ER remodeling is wholly dependent on a single viral gene product, UL148. Of course, there are other examples of individual viral gene products that are necessary to grossly perturb ER morphology during infection which are also found to be sufficient to induce such perturbations [e.g., (29, 30, 51, 52)]. Nonetheless, the ER structures caused by UL148 are— to the best of our knowledge, a hitherto undescribed ultrastructural characteristic of the HCMV infected cell, which is surprising given their size, scale, and prominence. Indeed, the average volume of the structures at 96 h postinfection amounts to roughly 60% of the size of the nucleus of an uninfected human fibroblast [∼500 µm^3^, (53)]. Although UL148 expression is accompanied by UPR activation, ER reorganization occurs without killing the host cell, even in the setting of ectopic expression when viral functions that inhibit programmed cell death are absent. Nor does UL148 affect the yield of infectious virus during replication in fibroblasts, since *UL148*-null mutant viruses replicate indistinguishably from wildtype parental virus in this cell type (1, 8). As with all enveloped viruses, HCMV requires the host cell secretory pathway to fold and process enormous quantities of viral envelope glycoproteins that endow progeny virions with the molecular machinery for infectivity. It is intriguing to consider how the infected cell tolerates spatial reorganization of ER membranes and of the associated glycoprotein quality control machinery without impacting the production of infectious progeny virions.

Furthermore, certain aspects of ER reorganization invoked by UL148 may be novel to cell biology. In particular, the densely packed collapsed ER in close association with varicosities that we observe to depend on UL148 during infection appear to differ from the ER structures induced by other ER perturbagens. For instance, a hereditary form of childhood cirrhosis caused by the *Z* allele of the alpha-1-antitrypsin (Z-A1AT) gene (*SERPINA*) is characterized by polymerization of Z-A1AT within the ER (26), leading to its accumulation in membrane-delimited inclusions (26, 27, 54). However, Z-A1AT inclusions are not observed to associate with regions of collapsed ER, nor does Z-A1AT suffice to activate the UPR, even though its expression sensitizes cells to other triggers of ER stress (27, 55).

HCMV encodes at least one other ER-resident immunevasin that activates the UPR, US11 (56). UL148 binds the NK-cell and T-cell costimulatory ligand CD58 to prevent its transport to the cell surface while US11 targets the heavy chain of class I major histocompatibility complex (MHC) for ER-associated degradation (57, 58). UL148 causes CD58 to accumulate as an immature glycoform, presumably within the ER, but does not lead to any obvious decrease in its overall abundance (9). Albeit that intracellular forms of CD58 are in our hands refractory to detection by standard indirect immunofluorescence protocols (data not shown), it seems reasonable to hypothesize that UL148 sequesters CD58 into the unusual ER structures that it induces. Moreover, our detergent solubility results suggest that UL148 may form aggregates or polymers in the course of preventing CD58 presentation at the cell surface (FIG 8). It is worth noting that UL148, an immunevasin targeting CD58, induces dramatic morphologic rearrangements of the ER, while another HCMV immunevasin, US11, targeting MHC class I for degradation evidently does not do so, even though both trigger ER stress. Nonetheless, it seems unlikely that retention of CD58 would be required for ER reorganization. In fact, the capacity of UL148 to retain CD58 may be functionally separable from its peculiar effects on the morphology and organization of the ER.

Despite that the relationship between the mechanism for CD58 retention and the formation of UL148-dependent ER structures remains unresolved, our findings may suggest a mechanism to explain the influence of UL148 on the tropism of the virus for epithelial cells. These effects, exemplified by a ∼100-fold replication advantage of *UL148*-null viruses during infection of epithelial cells, correlate with decreased expression of glycoprotein O (gO), a viral envelope glycoprotein, both in virions and in infected cells (1). We previously reported that gO behaves as a constitutive ERAD substrate during infection and that immature, newly synthesized forms of gO show enhanced stability in the presence of UL148 (8). Our findings herein show that UL148 causes large-scale sequestration of cellular factors important for ERAD, such as the ER mannosidase EDEM1 and the E3 ubiquitin ligase SEL1L/Hrd1, into large membranous structures. Moreover, the reticular ER, as indicated by the staining patterns of antibodies specific for calreticulin, PDI, and the KDEL motif, appears to remain largely intact following UL148-induced ER reorganization (FIG 3, 10). Since these observations indicate that UL148 depletes ERAD factors from the ER during the formation of the unusual structures, regions of ER not drawn into the structures might be expected to offer a more permissive folding environment for polypeptides, such as gO, which either fold slowly or inefficiently assemble into multiprotein complexes.

Based on these observations, one might hypothesize that UL148 alters ER proteostasis by physically dislocating (or sequestering) key elements of the “mannose removal time clock” system that marks poorly folding glycoproteins for destruction via ERAD (59, 60). Because gO is both the defining subunit of the heterotrimeric gH/gL/gO envelope glycoprotein complex that governs HCMV entry and cell tropism (5-7, 61, 62), and a constitutive ERAD substrate (1, 8), ER reorganization might be required for the effects of UL148 on HCMV cell tropism. Going forward, it will be crucial to determine whether classical ERAD substrates, such as the null Hong Kong variant of alpha-1-antitrypsin (63, 64) or ribophorin-332 (65, 66), are stabilized during UL148 expression, as would be predicted if UL148 shifts ER proteostasis to negatively modulate ERAD.

In addition to having found that factors involved in proteasomal ERAD, such as Hrd1 and EDEM1, are enriched at the UL148 structures, we observed that GABARAP, a mammalian ortholog of yeast ATG8, localizes to the UL148 ER structures during infection, and that another ATG8 ortholog, LC3B, and GABARAP both associate with the structures during ectopic expression of UL148. This may suggest roles for autophagy-related pathways in UL148-dependent reorganization of the organelle. Since our results indicate that UL148 is degraded by a proteasome-dependent pathway and not by lysosomes (**FIG S7**), it seems unlikely that selective autophagy of the ER is directly involved in recycling or degrading these ER structures. Nonetheless, ATG8 family proteins may contribute to formation of the globular ER structures, since we observed trafficking of UL148-GFP puncta to form large aggregates during live cell imaging (**FIG 12, 15, Movies S2, S8**). The literature indicates that misfolded proteins traffic in a microtubule (MT)-dependent manner to form pericentriolar structures termed aggresomes, which in the case of ERAD substrates such as the ΔF508 mutant of the cystic fibrosis transmembrane conductance regulator (CFTR), contain deglycosylated protein that presumably has already undergone dislocation from the ER (67, 68). ATG8 family proteins such as LC3B and GABARAP not only play roles in degradation of substrates via macroautophagy, but also bind MTs and recruit machinery for MT-dependent transport of cargos (69–71). Therefore, GABARAP and LC3B may be important for the recruitment of cellular machinery that transports perturbed ER cargoes to sites of large-scale accumulation.

Although our results argue that the ISR is required for ER reorganization during UL148 expression, precisely how UL148 triggers ER stress remains unknown. UL148 has been found to co-purify from cells with SEL1L, a component of the ERAD machinery. Thus, it seems plausible that UL148 may inhibit the Hrd1/SEL1L complex, which would cause the buildup of unfolded proteins and thus trigger the UPR. However, inhibition of ERAD in and of itself seems unlikely to account for the formation of ER structures. Another non-mutually exclusive possibility is that UL148 multimerizes or aggregates in a manner that constricts the ER lumen. For instance, the assembly of UL148 molecules on opposite sides of the ER lumen might constrict the organelle in a manner consistent with the collapsed regions of ER observed in our EM results (**FIG 6, 8**).

Additional work will be needed to decipher the molecular mechanisms by which UL148 causes reorganization of the ER, and to determine its physiological relevance in the setting of viral biology. Nonetheless, we have shown that UL148, when fused to a fluorescent protein (FP), suffices both to trigger and to indicate the presence of a functional ISR (**FIG 10-11, S5**). Hence, UL148-FP fusions may prove useful in high throughput chemical-genetic screens to identify novel small molecule inhibitors of the ISR as well as to identify cellular genes involved in stress-dependent remodeling of the ER. Interestingly, the two pharmacological agents that block UL148-dependent ER remodeling, ISRIB and GSK2606414, are known to prevent the formation of stress granules (SGs) (43, 72). SGs are comprised of condensed aggregates of protein and RNA which occur due to stalled mRNA translation in the context of disease states, such as amyotrophic lateral sclerosis (73), as well as during treatments with toxic agents such as arsenite (43, 72). It is fascinating to consider that mechanistic parallels exist between the formation of SGs and the UL148-dependent reorganization of the ER. Moreover, given the broad importance macromolecular aggregation in pathological conditions such as neurodegenerative diseases (74), certain of which also involve defects in ER proteostasis and aberrant activation of the UPR, UL148 may hold promise as tool to discover new agents to ameliorate disease.

## MATERIALS AND METHODS

### Cells and virus

hTERT-immortalized human foreskin fibroblasts (8), derived from ATCC HFF-1 cells (SCRC-1041) were maintained in Dulbecco’s modified Eagle’s medium supplemented with 5%-10% newborn calf serum (Millipore Sigma) and antibiotics (complete DMEM) exactly as described previously (8). i148^HA^ and i159^HA^ ARPE-19 epithelial cells (10), which upon treatment with 100 ng/mL doxycycline, express HA-tagged UL148 or Rh159, respectively, were likewise maintained in complete DMEM. For live cell imaging studies, i148^HA^ and i159^HA^ ARPE-19 were maintained in Opti-MEM (Thermo Fisher) supplemented with 3% certified tet-approved fetal bovine serum (FBS), 20 µg/mL gentamicin sulfate, 1 µg/mL puromycin HCl and 10 µg/mL ciprofloxacin HCl. Telomerase-immortalized rhesus fibroblasts (75) were a kind gift of Peter A. Barry, and were maintained in complete DMEM. The acute monocytic leukemia cell line THP-1 (TIB-202) was obtained from ATCC (Manassas, VA) and maintained as suspension cultures in RPMI 1640 medium supplemented with 10% FBS (Millipore Sigma) and antibiotics (10 µg/mL ciprofloxacin and 25 µg/mL gentamicin). THP-1 were differentiated into adherent macrophages by incubating for 48 h in the presence of 100 nM 2-O-tetradecanoylphorbol 13-acetate (Millipore Sigma), and subsequently infected with the indicated viruses at an MOI of 5 TCID_50_/cell.

Infectious bacterial artificial chromosome (BAC) clones of the following HCMV strains were used for this study: TB40/E (also known as TB40-BAC4 or TB_WT) (11) as well as its derivatives TB_148^HA^ and TB_148_STOP_, which were used in our previous studies (1, 8, 10); TR (TR*gfp*) (76, 77); Merlin repaired for *UL128* and *RL13* harboring *tetO* sequences upstream of *UL131* (pAL1393) (78); AD169 repaired for *UL131* (AD_r131)(1, 79) and AD_r131_148^HA^, a derivative of AD_r131 that carries a full length UL148 (from strain TB40/E) tagged with an HA-epitope at the original *UL148* locus (8). An infectious BAC clone of rhesus CMV (RhCMV) strain 68-1 (80) and a derivative that expresses an HA-tagged Rh159 (details below) were also used for certain experiments. The methods used for reconstitution of HCMV from purified BAC DNA, cultivation of virus and preparation of stocks, including ultracentrifugation through sorbitol cushions and determination of infectious titers by tissue culture infectious dose 50 (TCID_50_) assay have been described elsewhere (8, 10).

### Construction of recombinant viruses and new plasmids for the study

New recombinant viruses for this study were derived from BAC-cloned HCMV and RhCMVs using *en passant* mutagenesis in GS1783 *E. coli,* a strain K12 derivative that expresses the homing endonuclease I-SceI upon treatment with L-(+)-arabinose (81, 82). Recombinant BACs were confirmed by Sanger DNA sequencing of the modified regions, which was performed by Genewiz, Inc. (Piscataway, NJ) (not shown). Oligonucleotide primers for construction of recombinant viruses and plasmids were custom synthesized by Integrated DNA Technologies (Coralville, IA); sequences are provided in **Table S1**. Type II restriction endonucleases, T4 DNA ligase, and Gibson assembly reagents (NEB HiFi Assembly Master Mix) for generation of recombinant DNAs and for routine molecular cloning and subcloning procedures were obtained from New England Biolabs (Ipswitch, MA). KOD Hot Start DNA polymerase (EMD Millipore) was used for all PCR reactions.

To construct TB_159^HA^, a strain TB40/E derivative in which the UL148 open reading frame is replaced by Rh159 we carried out the following steps. Primers Rh159_Fw and Rh159_HA_Rv were used to amplify and HA-tag the *Rh159* ORF from pcDNA3.1+_Rh159_IRES_GFP (a gift from Klaus Früh, OHSU). The PCR product was inserted into EcoRV-digested pEF1*α* using Gibson assembly. A PCR product containing an ISce-I excisable kanamycin cassette was amplified from TB_148^HA^_*ISce-Kan* integrate BAC (1) using primers PpuISceKanGibs_Fw and PpuISceKanGibs _Rv and Gibson-assembled into PpuMI-digested pEF1*α*_Rh159 plasmid to yield plasmid pEF1*α*_Rh159_ISceKan. Primers Rh159_Fw_recomb and Rh159_Rv_recomb were used to generate a PCR product from template plasmid pEF1*α*_Rh159_ISceKan. The PCR product was electroporated into GS1783 E. coli harboring the TB_148^HA^ BAC, and Kan^r^ colonies harboring TB_*Δ*148_Rh159_ISceKan integrate BACs were isolated on Luria-Bertiani agar plates. Bacterial colonies representing kanamycin resistant integrates were obtained and were subsequently resolved to ‘scarlessly’ remove the positive selection marker by standard *en passant* protocols (81, 82) to yield TB_*Δ*148_Rh159^HA^, which we abbreviate herein as TB_159^HA^. TB_159^HA^ was sequence-confirmed using primers TB_159HAseq_F and TB_159HAseq_R. Similar strategies were used to modify the RhCMV 68-1 BAC (80) to incorporate sequences encoding an HA-epitope immediately before the stop codon of *Rh159*. An *en passant* strategy in which a shuttle plasmid carrying *I-SceI-Kan^r^* disrupted by in-frame nonsense codons was used insert premature nonsense codons in the *UL148* CDS in the context of HCMV strains Merlin (pAL1393) and ^TR^ (*TRgfp*), and was applied as described previously for generating TB_148_STOP_, the UL148_STOP_ mutant of strain TB40/E (8).

To construct “tet-on” lentivirus vector plasmids containing UL148 fused to GFP, we used primers Gibs_eGFP_Rv and 148_eGFP_Fw to amplify the GFP gene from a dsDNA gBlock, EGFP-P2A-3XHA, synthesized by Integrated DNA Technologies (Coralville, IA), which was a gift of Matthew D. Woolard (LSU Health Sciences Center, Shreveport, LA). In a separate PCR, the *UL148* gene was amplified from plasmid pcDNA3.1-UL148^HA^ (10) using primers Gib_148_Fw and 148_noStopRv. The two products were assembled, using Gibson Assembly, together with EcoRV opened pcDNA3.1(+) (Invitrogen) to produce pcDNA3.1-UL148-GFP. After confirming the absence of spurious mutations by Sanger DNA sequencing (Genewiz, not shown), the *UL148-GFP* cassette from pcDNA3.1-UL148-GFP was released by EcoRI and NotI digestion and ligated into pOUPc turboRFP-link plasmid which had been linearized using the same restriction sites. pOUPc turboRFP-link is a derivative of lentiviral vector pOUPc (10) that was constructed by reinsertion of turboRFP (tRFP). Briefly, tRFP was amplified from the original pOUPc-turboRFP using primer pair tRFP linker Fw and tRFP linker Rv and then reinserted by Gibson Assembly with pOUPc turboRFP that had been double-digested using EcoRI and Not I in order to linearize the vector and remove the *turboRFP* (RFP) sequence.

To express Rh159 fused to eGFP, primers Gibs_eGFP_Rv and 159_eGFP_Fw were used to amplify the GFP gene from EGFP-P2A-3XHA, and in a separate PCR reaction, primers Gibs_159_Fw and 159_noStop_Rv Rh159 were used to amplify *Rh159* from pcDNA3.1-Rh159^HA^ (10). The latter two PCR products were assembled together with EcoRV linearized pcDNA3.1(+) using Gibson Assembly, resulting in plasmid pcDNA3.1-Rh159. The latter plasmid was used as template in a PCR reaction with primers Gibs_Age_159_Fw and Gibs_Mlu_GFP_Rv. The resulting PCR product was Gibson assembled into the lentiviral vector pOUPc (10) which had been double-digested with Mlu I and Age I.

### Drug treatments

The PERK inhibitor, GSK2606414 (50), doxycycline hyclate, and folimycin were obtained from MilliporeSigma (Burlington, MA). ISRIB (42–46) was obtained from MilliporeSigma or APExBio (Boston, MA) and epoxomicin was procured Selleck Chemicals (Houston, TX). ISRIB was dissolved in DMSO to make a 10,000*×* stock solution (2 mM) and used at a final working concentration of 200 nM. GSK2606414 was dissolved in DMSO to make a 10,000*×* stock solution (11 mM) and used at a final working concentration of 1.1 µM. Folimycin was prepared as a 1000*×* (115.46 µM) stock solution in DMSO and used at 115 nM final. Epoxomycin was prepared as a 100*×* (2 mM) stock solution in DMSO and used at a final concentration of 20 µM.

### Confocal microscopy and live-cell imaging

Confocal indirect immunofluorescence microscopy imaging on fixed cells was carried out using Leica TCS SP5 Spectral Confocal Microscope (Leica Microsystems, Heidelberg, Germany) using a Leica HCX PL APO CS 63x/1.4-0.6NA objective under oil immersion, except for the image shown in Figure 1C, which was captured on a Nikon SIM-E and A1R confocal microscopy system (Nikon Instruments, Melville, NY) using a Nikon SR Apo TIRF 100x/1.49NA objective under oil immersion. Images for different fluorophore channels were acquired using sequential scanning. Direct immunofluorescence live cell imaging data were collected using the Nikon SIM-E microscope using a Nikon Apo 60x/1.40NA DIC objective. The 3D projection shown in FIG 1 was generated by NIS-Elements AR Analysis 4.60.00 (64-bit) software (Nikon) from Z-stacks captured on the Nikon SIM-E / A1R microscope using a Nikon SR Apo TIRF 100x/1.49NA objective. For FIG 11, 3D projections were generated by Imaris x64 9.3.0 software (Bitplane, Inc.) in maximum intensity projection (MIP) mode.

For fixed cell imaging experiments other than those shown in Fig S1D, cells were seeded on 12 mm circular, No. 1 thickness microscope cover glass (200121; Azer Scientific, Morgantown, PA). At the indicated times post-treatment and or post-infection, cells were washed with phosphate buffered saline (PBS) [137 mM NaCl, 2.7 mM KCl, 10 mM Na_2_HPO_4_, and 1.8 mM KH_2_PO_4_, pH 7.4]; fixed for 15 min at room temperature in PBS containing 4% (wt/vol) paraformaldehyde (Fisher Scientific, Waltham, MA), washed in PBS, permeabilized for 10 min using 0.1% Triton X-100 (in PBS), subsequently washed again in PBS, and then blocked for 45 min at 37 °C in PBS containing 5% (vol/vol) normal goat serum (Rockland Immunochemicals, Limerick, PA). Cells were then washed three times in PBS followed by incubation in 1% Human Fc Block (BD Biosciences, San Jose, CA) in PBS, for additional 45 min at 37 °C. Cells were then incubated in the presence of primary antibodies for 1 h at 37 °C or 4 °C overnight, and then washed three times with PBS containing 0.1% Tween-20 (PBST) for 5 min per wash. Alexa Fluor-labeled goat polyclonal secondary antibodies (all from ThermoFisher Invitrogen, Waltham, MA, see **Table S2**) were used for secondary detection. The slides were then mounted using Prolong Gold anti-fade reagent containing DAPI (ThermoFisher) and placed under a Leica TCS SP5 confocal microscope for image acquisition using a Leica 63*×* oil immersion objective lens (Leica Microsystems, Buffalo Grove, IL).

For indirect immunofluorescence staining results of primary clinical isolates (SI Fig S1D), four clinical HCMV isolates obtained by routine testing of throat swabs from patients of the Ulm University Medical Center were provided by the diagnostic laboratory of the Institute of Virology in Ulm. Sample material was applied to human fibroblasts and incubated for several days until HCMV-positive cells could be detected. Infected cells were then seeded together with uninfected fibroblasts, incubated for up to 5 days until plaques were formed and processed for indirect immunofluorescence staining. ERQC compartments were detected by staining for calnexin (CNX, E10; Santa Cruz Biotechnology, mouse, 1:50 dilution), the cVAC was detected by staining for HCMV pUL71 (rabbit, 1:500). Secondary antibody used for detection of CNX was Alexa Fluor 555 labeled goat anti-mouse IgG (1:1000) and for pUL71 detection, Alexa Fluor 488 conjugated goat anti-rabbit IgG (1:1000). Confocal images were acquired using the 63*×* objective of a Zeiss Observer Z1 fluorescence microscope equipped with Apotome and Zen software 2.3 (Carl Zeiss Microscopy GmbH, Jena, Germany).

### FRAP

FRAP studies were carried out on using live ARPE-19 cells from the i148^GFP^ and i159^GFP^ populations as follows. Cells were seeded as above for live-cell imaging, induced for transgene expression using dox (100 ng) for 24 h, and then placed on an incubated sample stage of a Nikon A1R SIM-E imaging system equipped with an SR Apo TIRF 100x/1.49NA objective (Nikon). Three rectangular regions in image fields were defined for measurement of (i) background signal, and of two regions with comparable initial GFP signals, (ii) one designated as a control region (no photobleaching) and another for (iii) photobleaching and recovery of signal after photobleaching. Photobleaching of selected regions was carried out for 20 s using 405 nm laser light from a LU-N4 laser fiber (Nikon) [power at source: 15 mW, power at objective: 8 mW]. Images and signal intensity measurements were captured every 2 s at a rate of 1 frame per s (488 nm excitation, FITC channel) starting immediately before photobleaching (t=1), and from t=20 s to t=320 s after.

### Electron microscopy

Procedures to prepare samples for transmission electron microscopy (TEM) included high-pressure freezing (HPF), freeze substitution, and Epon embedding, which were carried out as described previously (83). Briefly, human fibroblasts were seeded in µ-Slides (Ibidi GmbH, Martinsried, Germany) containing carbon-coated sapphire discs (Engineering Office M. Wohlwend GmbH) 1 day prior to infection at 80 to 90% confluence. Cells were infected with virus overnight at MOI 1. Medium containing viral inocula was replaced with fresh medium the next day. Infected cells on sapphire discs were fixed by using HPF with a Compact 01 high-pressure freezer (Engineering Office M. Wohlwend GmbH, Sennwald, Switzerland) at 5 dpi. Thereafter, cells on sapphire discs were processed by freeze-substitution and subsequently embedded in Epon (Fluka, Buchs, Switzerland). Ultrathin sections of the Epon-embedded cells were cut with an ultramicrotome (Ultracut UCT; Leica Microsystems, Wetzlar, Germany) and placed on Formvar-coated single-slot grids (Plano GmbH, Wetzlar, Germany). Grids were examined in a Jeol JEM-1400 (Jeol Ltd., Tokyo, Japan) transmission electron microscope equipped with a charge-coupled-device (CCD) camera at an acceleration voltage of 120 kV. Fixation and embedding of infected cells for scanning transmission electron microscopy (STEM) tomography was the same as described for TEM. Additional sample preparation steps were conducted as described previously (84, 85). Tomogram acquisition was conducted on a STEM Jeol JEM-2100F with an acceleration voltage of 200 kV. Tilt series were acquired from 600 nm thin sections from +72° to -72° with a 1.5° increment using the bright field detector. Image series were reconstructed to tomograms by weighted back projection with the IMOD software package (86). 3D visualization of the membrane structures was performed using Avizo lite software (Visualization Science Group, Burlington, MA, USA) by threshold segmentation.

### Western blotting

Western blotting procedures, including primary and secondary antibodies used for detection of HA tag, UL148, ATF4, eIF2*α*, P-eIF2*α* (Ser51), the HCMV viral nuclear antigen IE1, and the conditions used for detection of Ser51 phosphorylated eIF2*α*, were carried out as described previously (8, 10). Additional antibodies used in this study were mouse anti-GAPDH (Cat. No. 60004-1, Proteintech, Rosemont, IL,), mouse anti-gB clone 27-180 (87) (a generous gift of William J. Britt), and anti-GFP (D5.1) XP^®^ Rabbit mAb #2956 (Cell Signaling Technology, Danvers, MA).

### Solubility analyses

2.0 × 10^5^ human fibroblasts were infected at an MOI of 1 TCID_50_ per cell with TB_148^HA^ or TB_159^HA^. The following day, cells were washed twice with PBS to remove viral inoculum and replenished with DMEM containing 5% newborn calf serum. At the indicated times post-infection, cells were washed once with PBS and lysed by direct addition of 100 µL RIPA buffer [25 mM HEPES (pH 7.5), 400 mM NaCl, 0.1% SDS, 0.5% sodium deoxycholate, 1% NP-40, supplemented with 1*×* protease inhibitor cocktail (ApexBio)]. Lysates were collected and rotated at 4°C for 1 h. Insoluble material was pelleted by centrifugation (21,000 × *g*, 35 min). Supernatants containing soluble material were transferred to a fresh microfuge tube, and 33 µL of 4× Laemmli sample buffer [200 mM Tris (pH 6.8), 8% SDS, 40% glycerol, 0.08% bromophenol blue] was added to bring final volume to 133 µL. The pellet was disrupted in 133 µL of 1X Laemmli buffer prepared by diluting 4× Laemmli buffer in RIPA buffer. Samples were reduced by addition of beta-mercaptoethanol (5% final, v/v) and boiled at 90°C for 10 min. 40 µL of each sample was resolved by SDS-PAGE (12% polyacrylamide gel), transferred overnight to nitrocellulose membrane, and immunoblotted with antibodies against HA epitope or HCMV gB. Quantitation of secondary antibody fluorescence signals were performed using an Odyssey CLx scanner (Li-Cor, Inc., Lincoln, NE) in auto-scan mode. For each time-point, the signals from RIPA-soluble and insoluble bands were summed to yield total signal, and the ratio of insoluble band signal to total signal were also reported as percent insoluble HA over total HA signal.

### Statistical analyses

Statistical analyses were carried out using GraphPad Prism 8.1.0 for MacOS (GraphPad, Inc., San Diego, CA).

## ACKNOWLEDGEMENTS

This project was supported by NIH grants R01-AI116851 (to J.P.K.) and P30-GM110703. Its contents are solely the responsibility of the authors and do not necessarily represent the official views of the NIAID or the NIGMS.

We are especially grateful to Erik L. Snapp (HHMI, Janelia Laboratories) for helpful discussions and advice on interpretation of EM and confocal image data. We also thank the following individuals for generously sharing reagents: Dong Yu (Washington University, St. Louis, MO, currently GSK Vaccines), Thomas E. Shenk (Princeton University, Princeton, NJ), Richard J. Stanton (Cardiff University, United Kingdom), William J. Britt (University of Alabama, Birmingham), W. L. William Chang and Peter A. Barry (both of the University of California, Davis; Davis, CA), Christian Sinzger (University Medical Center Ulm, Ulm, Germany), Klaus Früh (Oregon Health Sciences University, Beaverton, OR), and Gregory A. Smith (Northwestern University, Chicago, IL).

## AUTHOR CONTRIBUTIONS

Performed experiments: HZ, CR, CCN, MNAS, JvE. Data analysis and interpretation: HZ, CR, CCN, JvE, CS, JPK. Designed experiments: HZ, CR, CCN, JvE, and JPK. Contributed new reagents: HZ, CH, CCN, JPK. Electron microscopy and STEM tomography: CR and JvE. Obtained funding: JPK. Wrote the manuscript: JPK with comments from JvE and CR.

---

**FIG S1:**
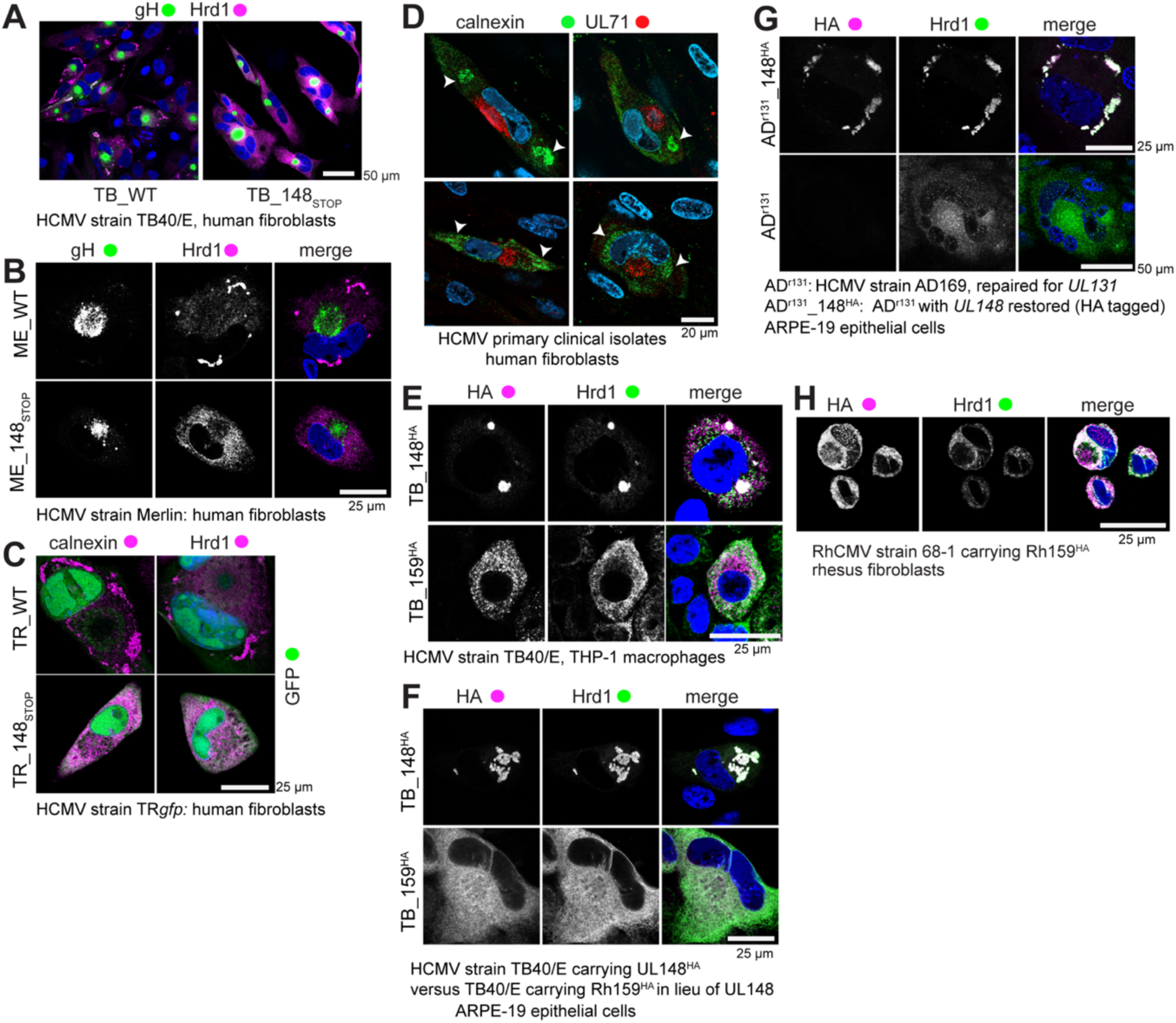
Additional confocal microscopy results from infected cells. (**A**) Lower magnification overview of results from FIG 1A-B; staining of gH and Hrd1 human fibroblasts 96 h postinfection (hpi) with wildtype HCMV strain TB40/E (WT) or *UL148*-null derivative TB_148_STOP_. (**B**) gH and Hrd1 in human fibroblasts 96 hpi with either wildtype strain Merlin recovered from BAC pAL1393 (ME_WT) or a *UL148*-null derivative of the same virus (ME_148_STOP_). (C): gH and Hrd1 at 96 hpi of human fibroblasts with wildtype HCMV strain TRgfp recovered from BAC-cloned TRgfp (TR_WT) or a *UL148*-null derivative of the same virus (TR_148_STOP_). (D) Staining of the viral tegument protein UL71 and calnexin in human fibroblasts 5 days postinfection with primary clinical HCMV isolates obtained from patient throat swabs. (**E**) HA and Hrd1 in THP-1 macrophages fixed 96 hpi with HCMV strain TB40/E derivative viruses TB_148^HA^ or TB_159^HA^. (**F**) HA and Hrd1 in ARPE-19 epithelial cells infected for 96 h with TB_148^HA^ or TB_159^HA^. (**G**) Staining of HA and Hrd1 in ARPE-19 infected for 96 h with HCMV strain AD169 repaired for *UL131* and to which an HA-tagged *UL148* from TB40/E was restored to the native *UL148* locus (AD^r131^_148^HA^) versus parental AD169 repaired for *UL131* (AD^r131^). Notes: (i) *UL131* is required for efficient infection of epithelial cells; (ii) for ADr131 we did not employ a viral marker (e.g., HA) to identify infected cells, so the appearance of syncytia and the characteristic kidney-bean shaped nucleus was used to indicate infected cells and a slightly lower magnification was used to best show these features. (**H**) HA and Hrd1 in telomerase-immortalized rhesus fibroblasts infected with BAC-derived rhesus CMV (RhCMV) strain 68-1 carrying an HA-tag at the C-terminus of Rh159.

**FIG S2:**
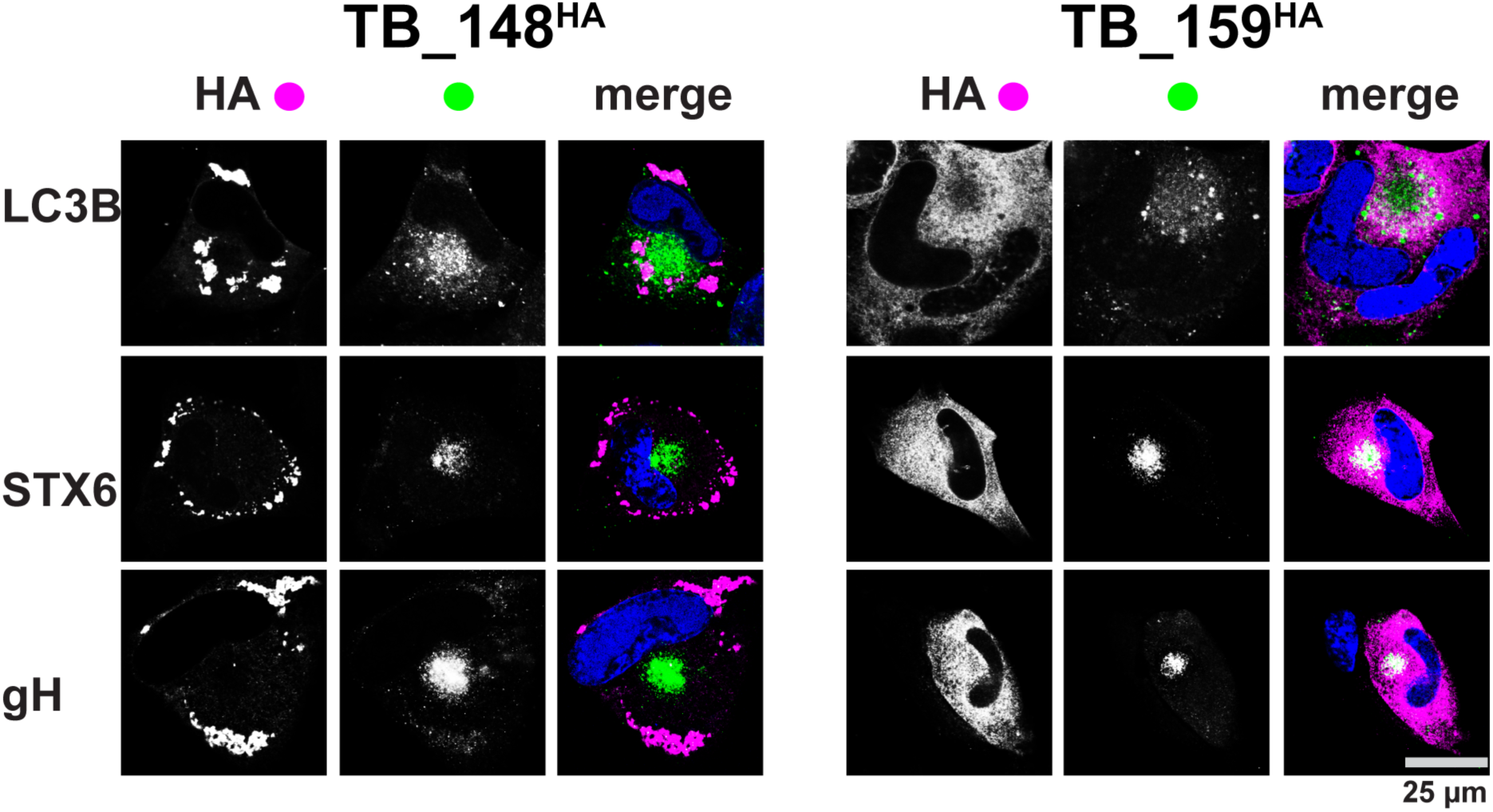
LC3B, syntaxin-6, and gH staining in HCMV infected cells. Human fibroblasts were infected with the indicated viruses for 96 h and then fixed, permeabilized, and stained using antibodies specific for the indicated proteins. Indirect immunofluorescence images were captured using a 63X objective on a Leica SP5 confocal microscope. gH: glycoprotein H, STX6: syntaxin-6.

**FIG S3:**
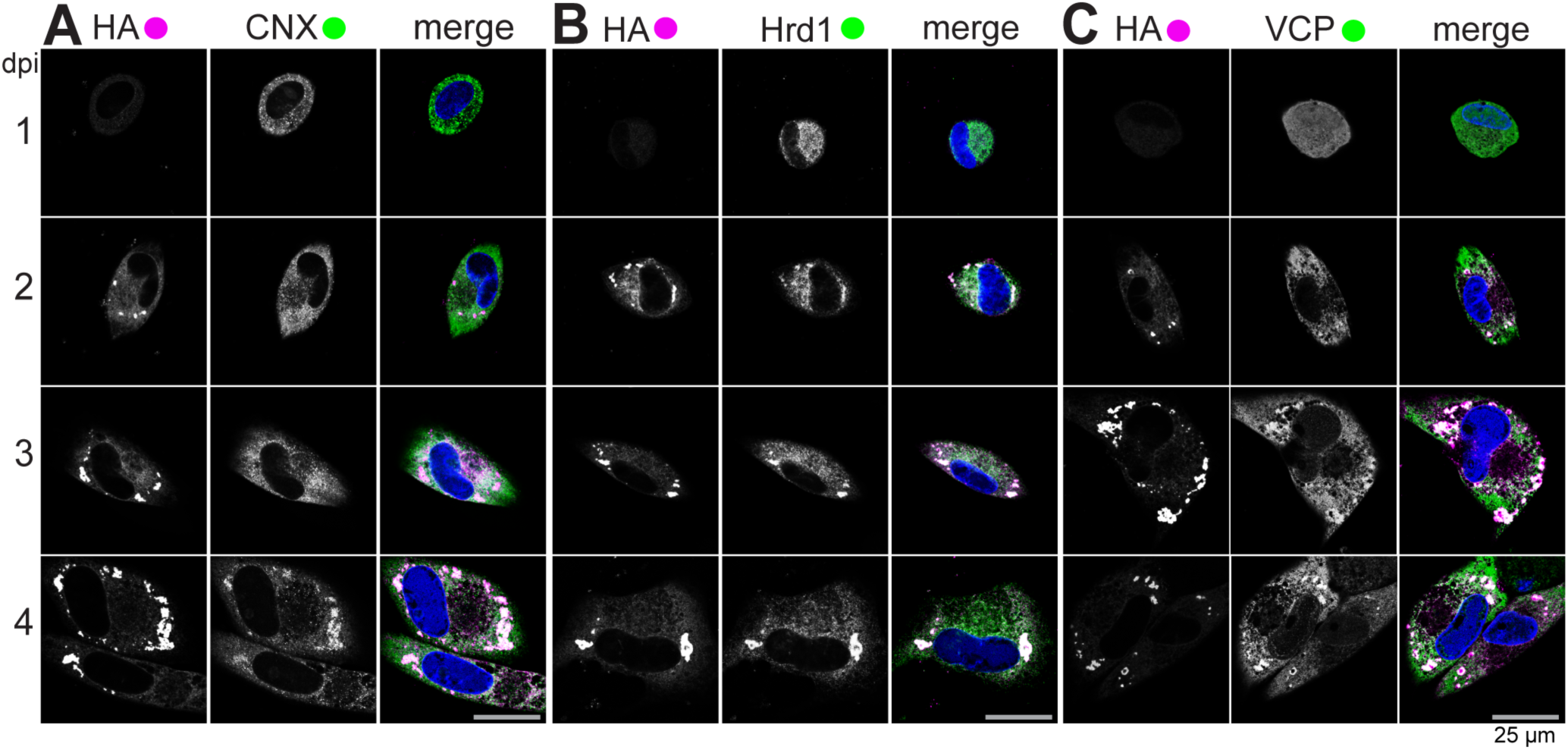
Cellular proteins involved in glycoprotein quality control are recruited with differing kinetics to UL148 ER structures. Fibroblasts infected with TB_148^HA^ at 1 MOI were fixed at the indicated time points (days post infection, dpi) and imaged by confocal microscopy after staining with antibodies specific for HA (UL148, magenta), CNX (green, **A**), Hrd1 (green, **B**) or VCP(green, **C**), DAPI (blue).

**FIG S4.**
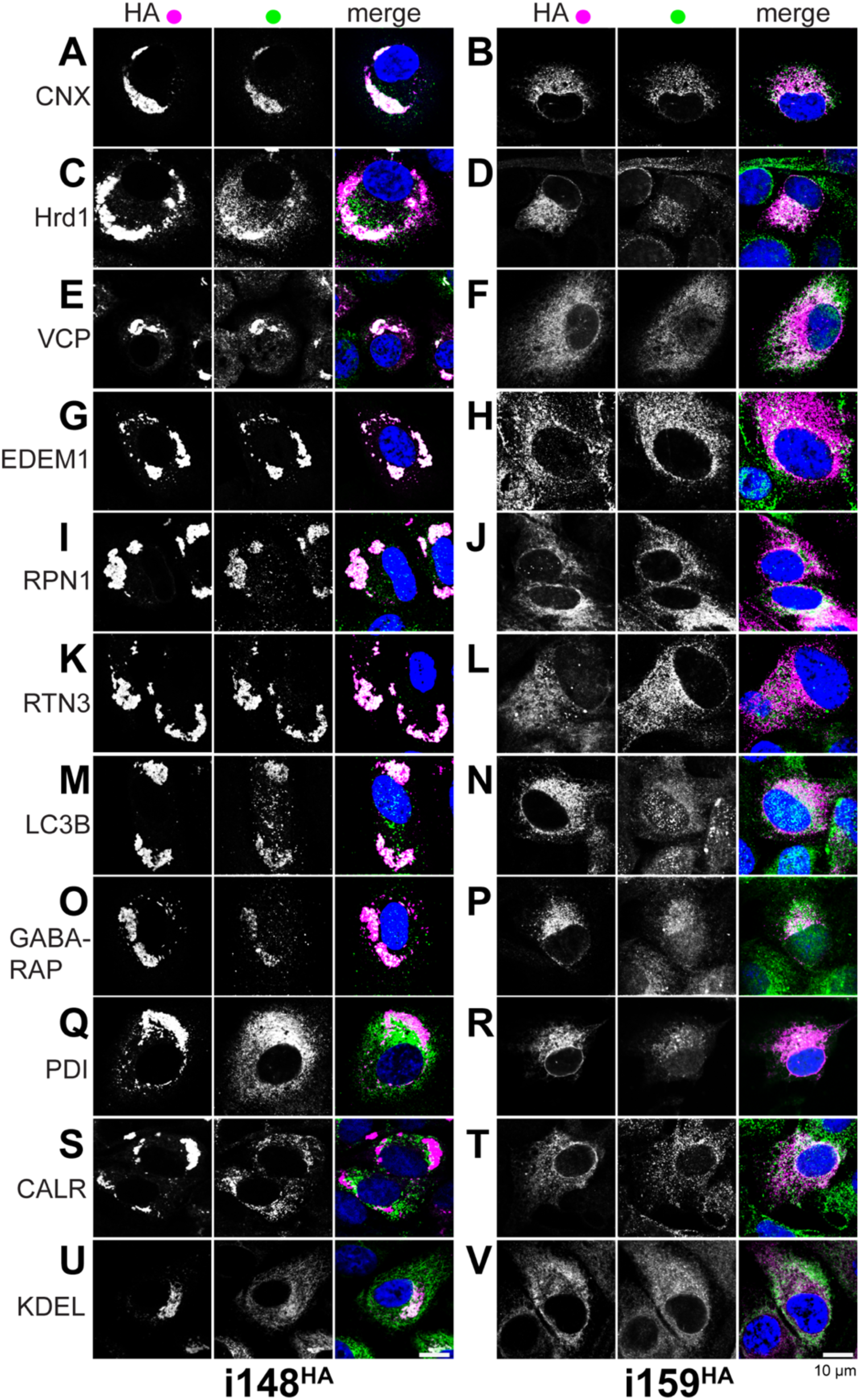
UL148 is sufficient to remodel the ER. Expression of HA-tagged UL148 or Rh159 was induced in “tet-on” ARPE-19 epithelial cells, i148^HA^ and i159^HA^, respectively. Cells were fixed at 48 h postinduction for indirect immunofluorescence staining for the indicated cellular markers (green) together with HA (magenta). Scale bar: 10 µm.

**FIG S5.**
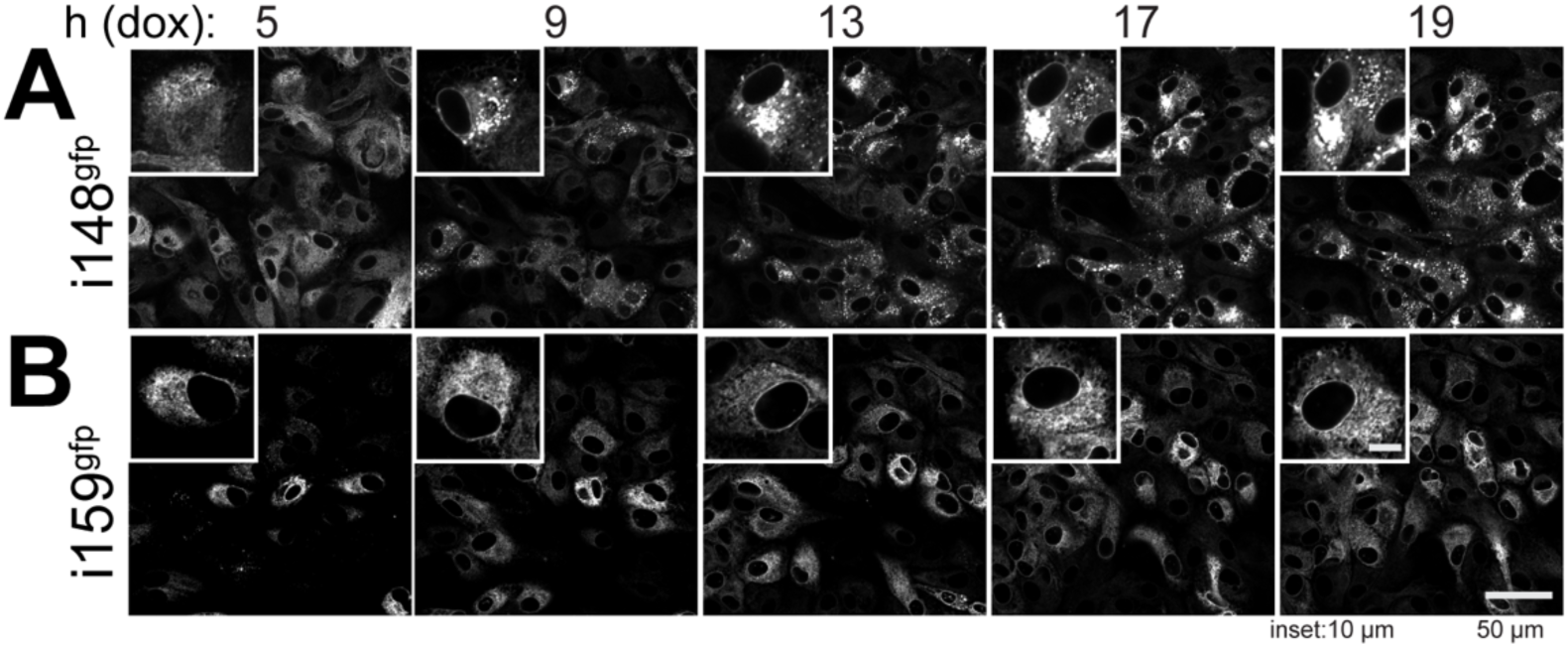
Live-cell Imaging of UL148-GFP and Rh159-GFP during induced ectopic expression. ‘Tet-on’ ARPE-19 epithelial cells that inducibly express either UL148 or Rh159 fused to green florescent protein (gfp), i148^gfp^ (**A**) and i159^gfp^ (**B**), respectively, were induced for transgene expression using 100 ng/mL doxycycline (dox) and imaged using live-cell microscopy. Images from the selected time points (h post treatment with dox, hpt) are shown. The main scale bar represents 50 µm. For inset panels at upper left of each image, which are magnified 2.4*×* relative to the main image, the scale bar represents 10 µm. Also see SI Movies S2-S3.

**FIG S6.**
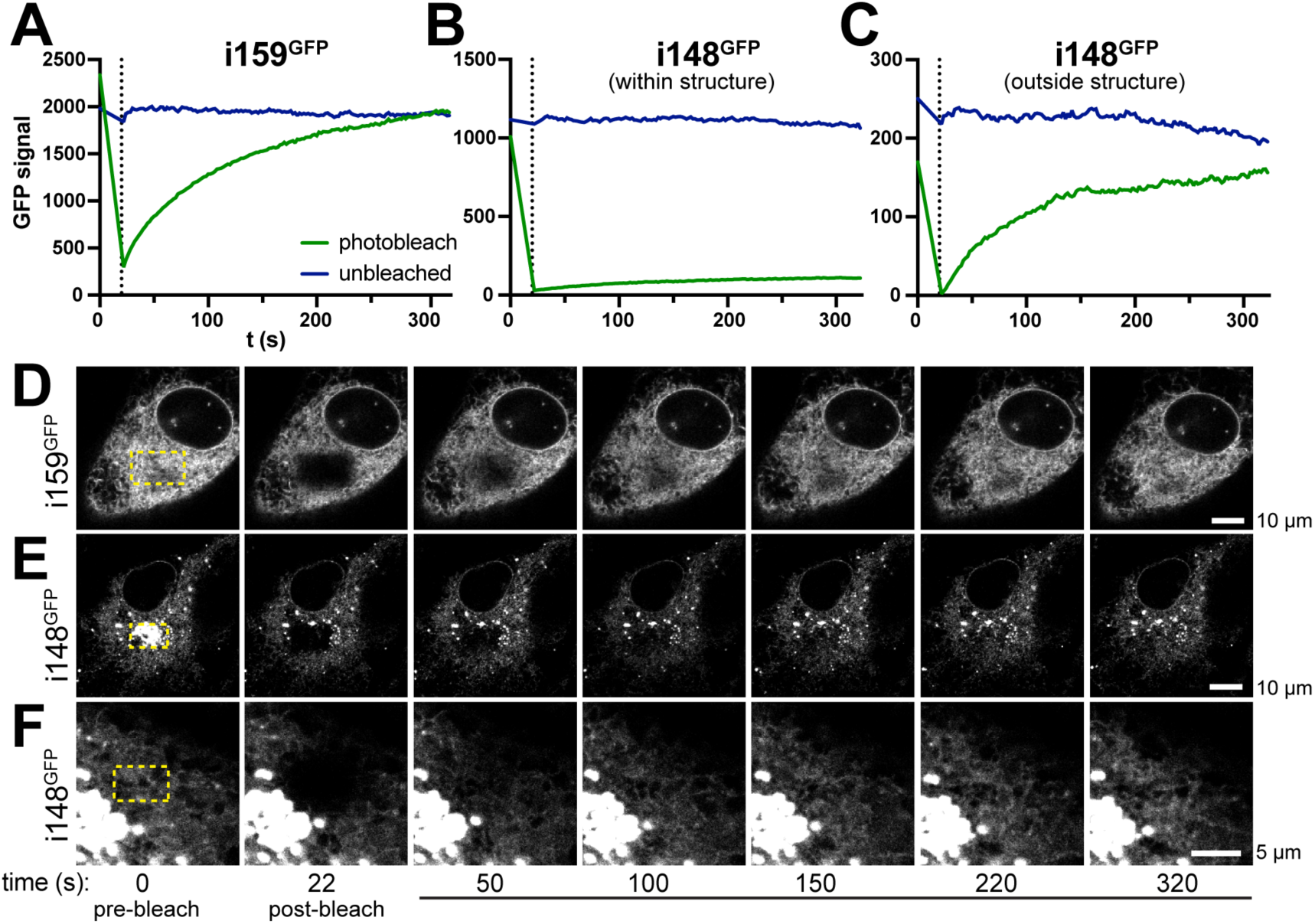
FRAP. ARPE-19 cells that inducibly express Rh159-GFP, i159^gfp^ (**A**) or UL148-GFP, i148^gfp^ (**B-C**) were doxycycline induced for 24 h (100 ng/mL doxycycline (dox). Separate regions of GFP signal in selected cells were photobleached (405 nm laser) or left unbleached, while a third region lacking GFP signal was chosen as a background reference were measured before and during fluorescence recovery after photobleaching (FRAP). Note: background signal was not plotted because values were resolvable from the x-axis. GFP signal intensity is plotted over a time period (seconds, s) starting with an exposure at t=0 (immediately before photobleaching), and including measurements taken every 2 s after photobleaching (0-20 s) until termination of the measurement series at t=322 s (300 s of FRAP). (**D-F**) Images from the selected time points (s, seconds) immediately before and after bleaching, and during fluorescence recovery period. Also see SI Movies S4-S6.

**FIG S7:**
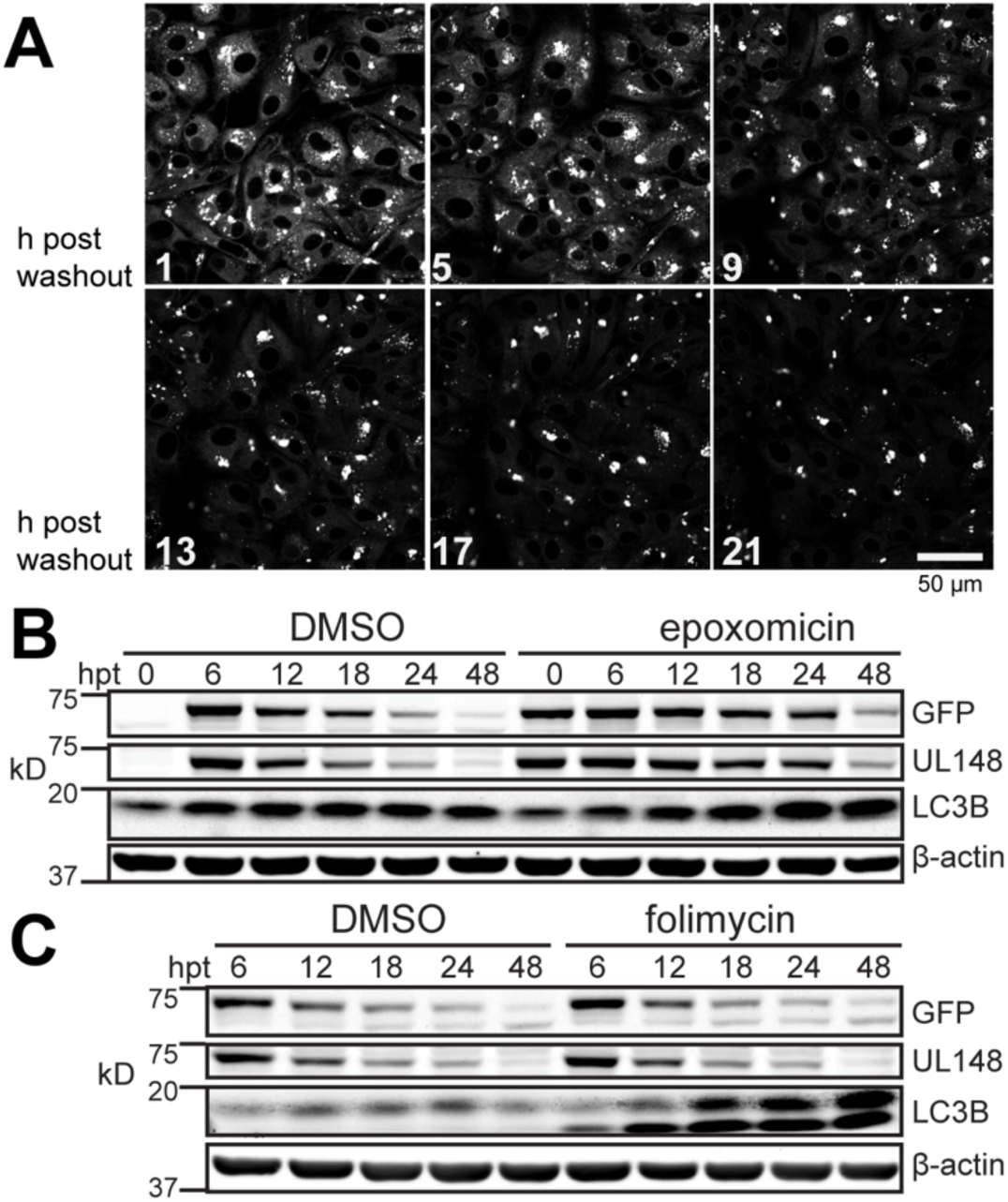
UL148-GFP is stabilized by inhibition of proteasomal but not lysosomal degradation. “Tet-on” ARPE-19 cells that inducibly express UL148-GFP (i148^gfp^) were induced for 24 h by the addition of 100 ng/mL doxycycline (dox), after which the dox inducing agent was washed out and medium containing either the proteasome inhibitor epoxomicin (20 µM) or the proton pump inhibitor folimycin (115 nM) was added and samples were harvested for Western blot analysis at the indicated times post treatment (h post treatment, hpt) with folimycin or epoxomicin. DMSO was added at 0.1% to control for folimycin, or 1% to control for epoxomicin. Also see SI Movie S7.

**TABLE S1.**
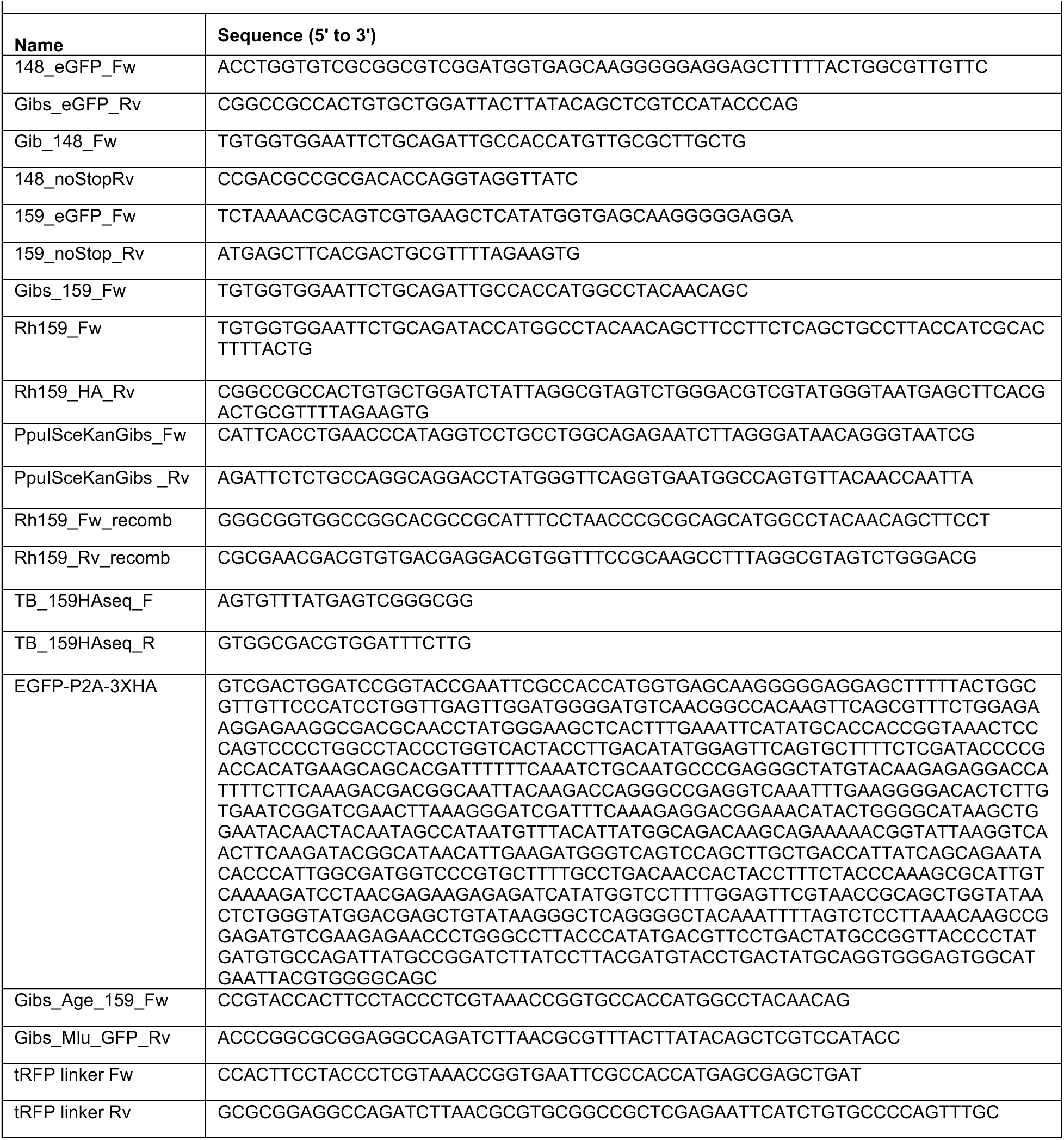
Oligonucleotide primers and synthetic DNAs.

**TABLE S2.** Antibodies Used in Immunofluorescence Microscopy.

| Primary Antibody | Host species | Isotype | Clone | Vendor | Catalog No. | Dilution used |
| --- | --- | --- | --- | --- | --- | --- |
| Calnexin | Mouse | mAb IgG1 | 37 | BD Biosciences | 610523 | 1:50 |
| Calnexin | Rabbit | mAb IgG | C5C9 | Cell Signaling Tech. | 2679 | 1:50 |
| Syntaxin 6 | Rabbit | mAb IgG | C34B2 | Cell Signaling Tech. | 2869 | 1:50 |
| SYVN1 (Hrd1) | Rabbit | mAb IgG | D3O2A | Cell Signaling Tech. | 14773 | 1:500 |
| SEL1L | Rabbit | pAb | n/a | Sigma | S3699 | 1:50 |
| KDEL | Mouse | IgG2a | 10C3 | Enzo Life Sci., Inc. | ADI-SPA-827-D | 1:50 |
| PDI | Rabbit | mAb IgG | C81H6 | Cell Signaling Tech. | 3501 | 1:50 |
| Herp (HerpUD1) | Rabbit | pAb IgG | n/a | Proteintech | 10813-1-AP | 1:50 |
| VCP | Mouse | IgG2a | 5 | Invitrogen / Life Technologies | MA3-004 | 1:50 |
| Calreticulin | Rabbit | mAb IgG | D3E6 | Cell Signaling Tech. | 12238 | 1:200 |
| Ribophorin 1 (RPN1) | Rabbit | pAb IgG |  | ThermoFisher / Invitrogen | PA5-65662 | 1:50 |
| EDEM-1 | Mouse | mAb IgM | D-1 | Santa Cruz | sc-377394 | 1:50 |
| GABARAP | Rabbit | mAb | E1J4E | Cell Signaling Tech. | 13733 | 1:200 |
| LC3B | Rabbit | mAb | D11 | Cell Signaling Tech. | 3868 | 1:200 |
| HA epitope | Chicken | IgY |  | Bethyl | A190-106A | 1:200 |
| HCMV glycoprotein H (gH) | Mouse | IgG2b | 14-4b | Gift of William J. Britt, M.D. | n/a | 1: 50 |
| Reticulon-3 (RTN3) | Mouse | IgG2a |  | ThermoFisher / Invitrogen | MA5-15538 | 1: 200 |
| HA epitope | Rabbit | pAb |  | Bethyl | A190-108A | 1:1000 |
| Secondary Antibody | Host species | Isotype |  | Vendor | Catalog No. | Dilution used |
| Alexa Fluor 488 anti-Mouse IgG (H+L), cross adsorbed | Goat | IgG |  | ThermoFisher / Invitrogen | A11001 | 1:1000 |
| Alexa Fluor 488 anti-Rabbit IgG (H+L) Cross-Adsorbed | Goat | IgG |  | ThermoFisher / Invitrogen | A11008 | 1:1000 |
| Alexa Fluor 594 anti-Rabbit IgG (H+L) Cross-Adsorbed | Goat | IgG |  | ThermoFisher / Invitrogen | A11012 | 1:1000 |
| Alexa Fluor 594 anti-Mouse IgG (H+L) Cross-Adsorbed | Goat | IgG |  | ThermoFisher / Invitrogen | A11005 | 1:1000 |
| Alexa Fluor 647 anti-Chicken IgY (H+L) | Goat | IgG |  | ThermoFisher / Invitrogen | A11039 | 1:1000 |

## SUPPLEMENTARY VIDEO FILES

**Movie S1:** STEM tomography of UL148 ER structures in wildtype HCMV infected fibroblasts at 5 dpi.

**Movie S2:** i148^gfp^ cells from 2-19 h post dox induction. Movie S3: i159^gfp^ cells from 2-19 h post dox induction. Movie S4: FRAP of UL148-GFP signal within ER structure. Movie S5: FRAP of UL148-GFP signal outside ER structure. Movie S6: FRAP of Rh159-GFP signal.

**Movie S7:** i148^gfp^ cells from 30 min to 22 h 07 min post washout of doxycycline (dox).

**Movie S8:** i148^gfp^ cells in the presence of 0.01% DMSO, from 2-19 h post dox induction.

**Movie S9:** i148^gfp^ cells in the presence of 1.1 µM GSK2606414, from 2-19 h post dox induction.

**Movie S10:** i148^gfp^ cells in the presence of 200 nM ISRIB, from 2-19 h post dox induction.

